# The CDKL5 kinase undergoes liquid-liquid phase separation driven by a serine-rich C-terminal region and impaired by neurodevelopmental disease-related truncations

**DOI:** 10.1101/2024.11.18.624084

**Authors:** Marco Dell’Oca, Stefania Boggio Bozzo, Serena Vaglietti, Antonia Gurgone, Vita Cardinale, Gregorio Ragazzini, Andrea Alessandrini, Luca Colnaghi, Mirella Ghirardi, Maurizio Giustetto, Ferdinando Fiumara

## Abstract

Mutations of the *cyclin-dependent kinase-like 5* (CDKL5) gene, which encodes a serine/threonine protein kinase, can cause the *CDKL5 deficiency disorder* (CDD), a severe neurodevelopmental disease characterized by epileptic encephalopathy and neurocognitive impairment. The CDKL5 kinase consists of a catalytic N-terminal domain (NTD) and a less characterized C-terminal domain (CTD). Numerous disease-related mutations truncate CDKL5, leaving the NTD intact while variably shortening the CTD, which highlights the importance of the CTD for CDKL5 function. By systematically analyzing CDKL5 compositional features and evolutionary dynamics, we found that the CTD is a low-complexity region (LCR) highly enriched in serine residues and with a high propensity to undergo liquid-liquid phase separation (LLPS), a biophysical process of condensation controlling protein localization and function. Using a combination of super-resolution imaging, electron microscopy, and molecular and cellular approaches, including optogenetic LLPS induction, we discovered that CDKL5 undergoes LLPS, predominantly driven by its CTD, forming membraneless condensates in neuronal and non-neuronal cells. A CTD internal fragment (CTIF) plays a pivotal LLPS-promoting role, along with the distal portion of the protein. Indeed, two disease-related truncating mutations (S726X and R781X), eliding variable portions of the CTIF, significantly impair LLPS. This impairment is paralleled at the functional level by a reduction in the CDKL5-dependent phosphorylation of EB2, a known CDKL5 target. These findings demonstrate that CDKL5 undergoes LLPS, driven by a CTD region elided by most disease-related truncating mutations. Its loss––through the impairment of CDKL5 LLPS and functional activity––may play a key role in the molecular pathogenesis of CDD.

## INTRODUCTION

Mutations in the *cyclin-dependent kinase-like 5* (CDKL5) gene, encoding a serine/threonine protein kinase, can cause the *CDKL5 deficiency disorder* (CDD), a neurodevelopmental epileptic encephalopathy characterized by early-onset seizures, severe neurocognitive impairment, and sensorimotor alterations, which was originally classified as a Rett syndrome variant (Leonard et al., 2022). Moreover, CDKL5 mutations can also lead to related neurodevelopmental conditions, like West syndrome, also characterized by epileptic encephalopathy (Kato, 2006).

The biological activity of the CDKL5 protein has been linked to the regulation of a variety of cellular processes ranging from DNA repair to synapse assembly and function in neurons (La Montanara et al., 2015; Canning et al., 2018; Khanam et al., 2021; Pizzo et al., 2016, 2020; Gurgone et al., 2023). However, only relatively few of its direct protein targets and mechanisms of action have been identified (Katayama et al., 2020; Van Bergen et al., 2022). Different isoforms of the CDKL5 kinase are generated by alternative splicing, displaying tissue-specific expression patterns, and one of these isoforms is highly expressed in the nervous system (Hector et al., 2016; Frasca et al., 2022). The domain architecture of the protein consists of an N-terminal domain (NTD) with catalytic function followed by a long C-terminal domain (CTD). While the structure and function of the catalytic domain are quite well characterized (Canning et al., 2018), those of the CTD remain largely elusive. The CTD, which bears two nuclear localization signals (NLSs) and a nuclear export signal (NES), is thought to regulate the subcellular localization and catalytic activity of CDKL5 (Bertani et al., 2006; Rusconi et al., 2008; Frasca et al., 2022).

Most CDD-related missense mutations affect the NTD and can impair or abolish its kinase domain function (Canning et al., 2018). Remarkably, however, numerous CDD-related frameshift/nonsense mutations, leading to the production of truncated variants of the protein, occur both in the kinase domain and in the CTD (Fehr et al., 2015; Leonard et al., 2022). Thus, numerous pathogenic forms of the protein bear an intact catalytic kinase domain and a variably shortened CTD. These findings indicate that the integrity of the kinase domain alone is not sufficient to prevent CDD, highlighting the fundamental importance of the CTD in ensuring the proper physiological function of the CDKL5 protein.

The clinical spectrum of CDKL5-related disorders is quite heterogeneous, with limited genotype-phenotype correlations linking specific mutations to defined clinical outcomes (Fehr et al., 2015; Demarest et al., 2019; Leonard et al., 2022). Thus, a detailed understanding of the molecular and functional features of the CDKL5 CTD may provide key knowledge to better understand both the physiological roles of this conspicuous protein domain, which accounts for ∼2/3 of the protein length, and the pathological impact of those CDD-related truncating mutations that variably shorten its length.

To address these issues, we performed a systematic *in silico* analysis of the compositional features, structural organization, and evolutionary dynamics of CDKL5. This analysis revealed that the CTD is a low-complexity region (LCR) of the protein highly enriched in serine residues and with a high predicted propensity to undergo liquid-liquid phase separation (LLPS), a biophysical process of condensation controlling protein localization and function. We found that CDKL5 is indeed able to form LLPS-driven condensates in neuronal and non-neuronal cells, as revealed by morphological, molecular dynamics, and biochemical criteria. Through a molecular dissection approach, we found that CDKL5 LLPS relies on the interplay between the CTD and the NTD, and that it is promoted particularly by a serine-rich CTD internal fragment (CTIF) together with the more distal portion of the protein. Two disease-related truncating mutations affecting this domain (i.e. S726X and R781X) impair both the LLPS behavior and the catalytic activity of CDKL5. These findings have important implications for our understanding of CDKL5 physiological function and reveal novel LLPS-related molecular underpinnings of CDD.

## RESULTS

### The CDKL5 CTD is highly enriched in serine residues

To gain a deeper understanding of the possible biological roles of the CDKL5 CTD, we undertook a systematic *in silico* analysis of its compositional and structural features in comparison with those of the whole CDKL5 protein, of its NTD, and of the entire human proteome.

We first analyzed the occurrence and distribution of the 20 amino acids in CDKL5 (the 960-residue isoform enriched in the brain, or CDKL5_107_; Frasca et al., 2022) and its two major domains, i.e. the NTD (a.a. 1-297), mostly occupied by the catalytic kinase domain (a.a. 13-297), and the downstream CTD (a.a. 298-960). Moreover, we quantified the mean occurrence of the 20 amino acids across all proteins in the Uniprot reference human proteome to identify over- and under-represented residues in CDKL5 and its two major domains (**Fig. 1A-C; Suppl. Fig. 1A,C**).

**Figure 1.**
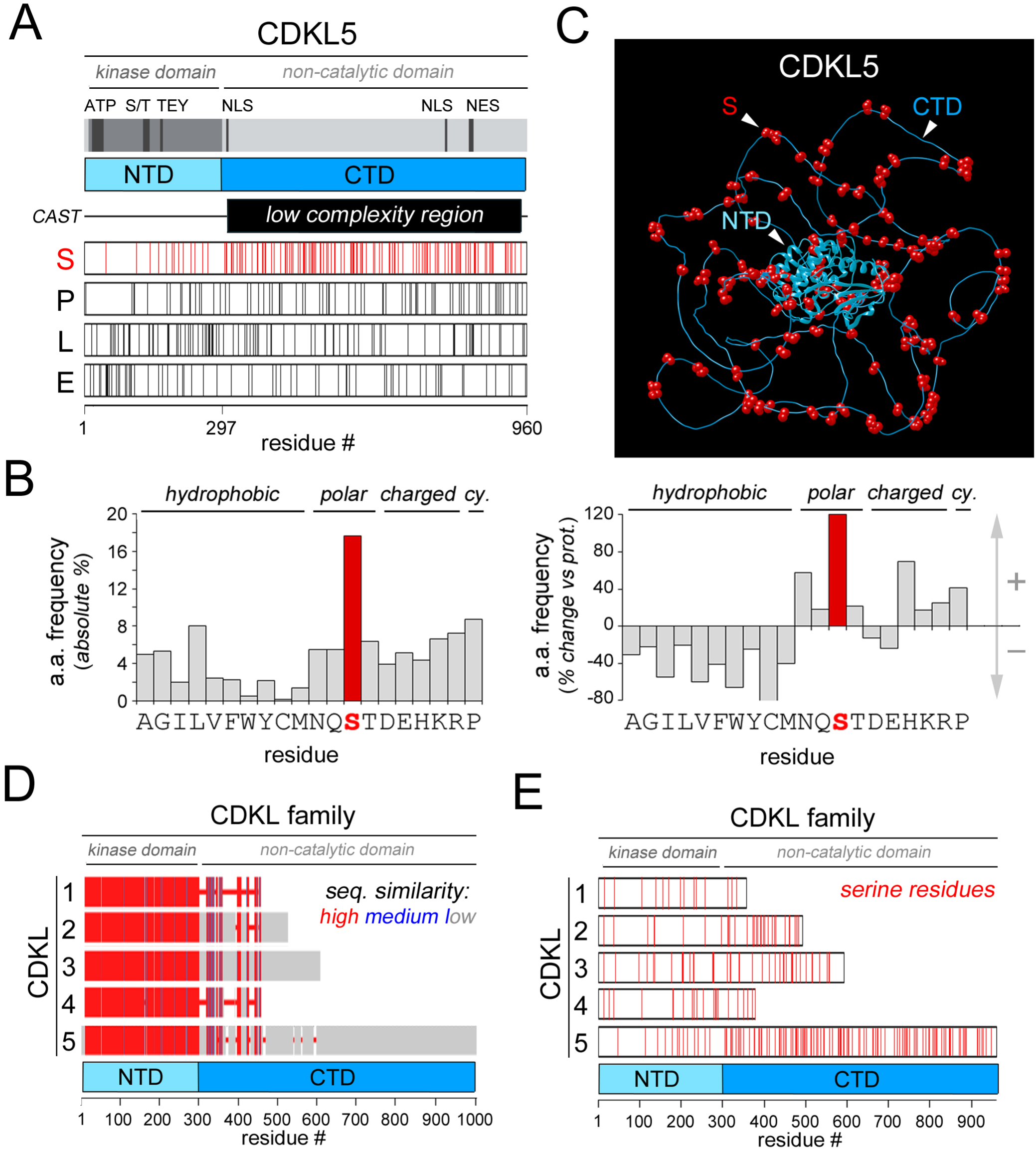
The CDKL5 CTD is highly enriched in serine residues. **A**. The *gray ba*r is a schematic representation of the CDKL5 protein highlighting the kinase domain (*dark gray*) and, in *black*, notable functional sites, i.e. the ATP-binding site (ATP), the catalytic site (S/T), and the activation loop site (TEY), two nuclear localization signals (NLS), and a nuclear export signal (NES). The *colored bar* highlights the N-terminal domain (NTD, *cyan*), containing the kinase domain, and the C-terminal domain (CTD, *turquoise*) formed by a low-complexity region, as identified by the CAST algorithm (*black bar*). *Thin vertical lines* in the four white bars below indicate the position of the indicated amino acid residues (S in *red*, P, L, and E in *black*). **B**. The bar graph on the *left* plots the percent occurrence of the 20 amino acids within the CDKL5 CTD. Note the high percentage of serine residues (*red bar*). The bar graph on the *right* plots the difference between the percent occurrence of the 20 amino acids in the CTD and their mean percent occurrence across all proteins of the human proteome. This difference reaches the highest value (∼120%) for serine residues. **C**. AlphaFold structural model of human CDKL5. The NTD (*cyan*), the CTD (*turquoise*), and serine residues (*red spheres*) are highlighted. The CTD displays a mostly random coil conformation, forming a shell around the central catalytic domain, and is highly enriched in serine residues. **D**. Schematic representation of an alignment of the primary sequences of human CDKL1-5 obtained using Cobalt (NCBI). *Red*, *blue*, and *gray* bars indicate, respectively, high, intermediate, and low degrees of sequence similarity along the primary sequence alignment. **E.** Schematic representation of CDKL1-5 (*white bars*) in which the position of serine residues is highlighted in *red*. Note how CDKL5 has the longest serine-rich CTD.

This analysis revealed that serine is the most represented amino acid, both in CDKL5 *in toto* (13.56%) and in its CTD (17.67%), largely exceeding (+68% and +119%, respectively) the mean percent occurrence of this residue in the human proteome (8.03%; **Fig. 1A,B**; **Suppl. Fig. 1A,B**). Consistent with these observations, the CAST algorithm identified the CTD overall as a low complexity region within CDKL5 (**Fig. 1A**). Notably, serine is much less represented in the catalytic NTD (4.39%, i.e. -45.35% *vs* proteome). Leucine (L) is instead the most represented amino acid (13.51%) in the NTD, together with charged glutamate (E; 9.8%) and lysine (K; 8.11%) residues (**Fig. 1A,B; Suppl. Fig. 1A,B**). When we analyzed the distribution of serine and leucine in CDKL5 (**Fig. 1A, Suppl. Fig. 1A**), we observed that serine residues are mostly concentrated in the CTD, scattered along its entire length, also in doublets or triplets. Leucine residues are instead more concentrated in the NTD, especially near the junction with the CTD, while the frequency of charged residues peaked in the initial portion of the NTD (**Fig. 1A; Suppl. Fig. 1A**).

The amino acid composition of the CTD appears to deviate substantially from the average composition of the human proteins. Indeed, the CTD appears to be enriched in polar (S,N,T,Q), positively charged (K,R,H) and proline (P) residues, and depleted of hydrophobic and negatively charged ones (**Fig. 1B**, **Suppl. Fig. 1B**).

These differential amino acid distributions across the two domains appear to mark structurally distinct regions of the protein. Indeed, as visible in the AlphaFold CDKL5 model (**Fig. 1C**), the serine-rich portion of the protein is predicted to form long random coil loops surrounding the central catalytic core like a shell. Leucine residues, by contrast, are concentrated within the α-helices of the kinase domain in the NTD, as shown by its experimentally determined structure in the PDB database (**Suppl. Fig. 1C**).

Such a serine-rich CTD is either absent or considerably shorter in the other four members of the human CDKL family of kinases (CDKL1-4), which display instead a high degree of sequence similarity with CDKL5 in the NTD (**Fig. 1D**). The amino acid composition of the five members of the CDKL family reflected this fact. Indeed, the occurrence of serine residues in CDKL1-4 is similar to that in the whole proteome (8.03%), or even lower, and is substantially higher only in CDKL5 (4.2% in CDKL1, 6.28% in CDKL2, 7.09% in CDKL3, and 5.54% in CDKL4, 13.54% in CDKL5, **Fig. 1E)**.

These findings indicate that the human CDKL5 CTD is a long LCR region, highly enriched in serine residues, that is unique to CDKL5 among its CDKL paralogs.

### The serine-richness of the CTD is phylogenetically conserved in vertebrates and peaks in mammals

To determine whether the observed enrichment in serine residues of the CDKL5 CTD is a human-specific or a phylogenetically conserved feature of the protein, we systematically analyzed the primary sequence composition of CDKL5 orthologs from 332 species belonging to major clades/taxa of different stem age along the vertebrate lineage (**Fig. 2A**). We found that the frequency of serine residues in the CTD is higher than that found in the NTD even in species belonging to clades/taxa with the oldest stem age, such as Agnatha (10.69% in *Petromyzon marinus*), with a progressive increase going towards younger clades, reaching its maximum in mammals (17.39% on average; **Fig. 2B-E**). Overall, the mean percent occurrence of serine residues in the CTD is significantly higher in mammalian than in non-mammalian species (17.39 ± 0.04, n = 133 species, *versus* 16.20 ± 0.08, n = 199, p<0.001, *t*-test; **Fig. 2D**). Moreover, the increase in serine occurrence in the CTD correlates significantly with clade/taxa divergence times along the vertebrate lineage (r = 0.81, n = 14, p<0.001; **Fig. 2E**).

**Figure 2.**
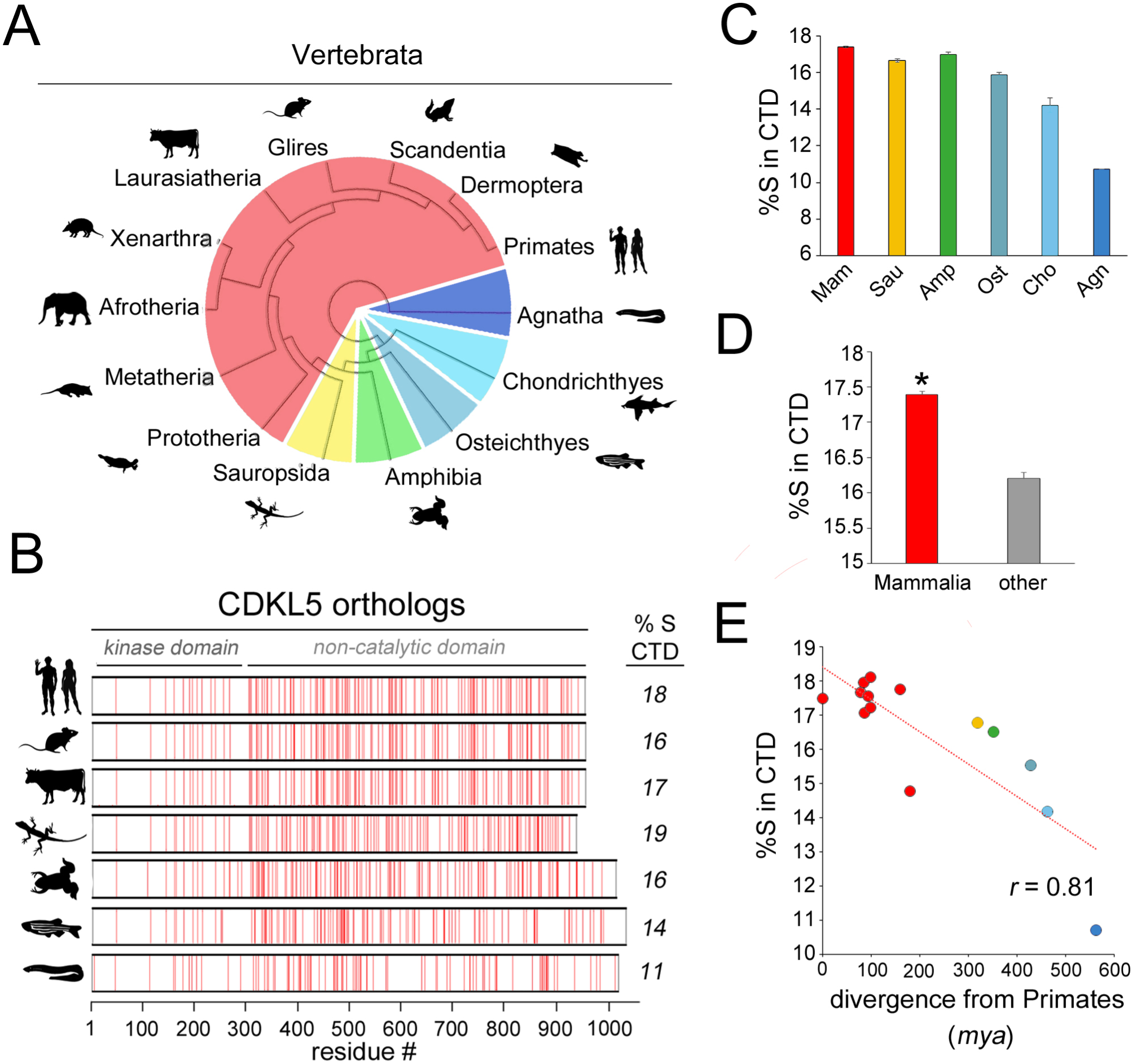
Evolutionary trends in the serine enrichment of the CDKL5 CTD in vertebrates. **A**. Phylogenetic tree encompassing some of the main clades/taxa of the vertebrate lineage. Mammalia, Sauropsida, Amphibia, Osteichtyes Chondrichthyes, and Agnatha are highlighted, respectively, in *red*, *yellow*, *green*, *teal*, *cyan*, and *blue*. **B.** Schematic representation of CDKL5 orthologs (*white bars*) of seven vertebrate species from representative vertebrate clades/taxa in which the position of serine residues is highlighted in *red*. The black silhouettes indicate, from *top* to *bottom*, *Homo sapiens*, *Mus musculus*, *Bos taurus*, *Anolis carolinensis*, *Xenopus laevis*, *Danio rerio*, and *Petromyzon marinus*. The enrichment in serine residues increases going from species belonging to clades/taxa with older stem age to species belonging to younger clades along the vertebrate lineage. **C.** Bar graph showing the mean percent occurrence of serine residues in the CTD of CDKL5 orthologs of the indicated clades/taxa (Mammalia, *Mam*, n = 133; Sauropsida, *Sau*, n = 108; Amphibia, *Amp*, n = 9; Osteichtyes, *Ost*, n = 79; Chondrichthyes, *Cho*, n = 2; and Agnatha, *Agn*, n = 1). Note the increase in serine occurrence going from Agnatha to Mammalia. **D.** Bar graph displaying the mean percent occurrence of serine residues in the CTD of CDKL5 orthologs from mammalian (*red*) *versus* non-mammalian species (*other*, *gray*). **E.** Scatterplot displaying the significant correlation between clade/taxa divergence times and the mean percent occurrence of serine residues in the CTD (n = 14 clades/taxa, as listed in Fig. 1A).

Taken together, these findings indicate that the enrichment in serine residues is an ancient feature of CDKL5 that has been then preserved, and even enhanced, in younger clades along the vertebrate lineage, strongly suggesting its functional relevance.

CDKL genes have also been identified in invertebrates, such as *Caenorhhabditis elegans* and *Drosophila melanogaster*. These species possess only a single CDKL gene, more closely related to the vertebrate CDKL1-4 group, bearing a relatively short CTD. Interestingly, the CTDs of both these two invertebrate CDKLs contain LCRs enriched in either alanine/glutamine (A/Q in *D. melanogaster*) or asparagine/serine (N/S in *C. elegans*), with an overall configuration similar to that of human CDKL4 (**Suppl. Fig. 2A)**. These observations suggest the possibility that LCRs may have appeared multiple times in CDKL genes through convergent molecular evolution (see *Discussion*).

### The serine-rich CTD is predicted to undergo liquid-liquid phase separation (LLPS)

As serine-rich LCRs are candidate mediators of protein oligo-/poly-merization and phase separation (Lilliu et al., 2018; Sabari et al., 2018), we sought to determine whether the CTD may promote CDKL5 LLPS. The FuzDrop algorithm predicted a high overall probability for the protein to undergo LLPS (*probability of spontaneous liquid-liquid phase separation*: (pLLPS) >0.99, on a 0-to-1 scale). Strikingly, most of the residues in the serine-rich CTD, unlike those in the kinase domain, displayed above threshold per-residue P_DP_ scores (≥ 0.6, on a 0-1 scale with a positive prediction threshold of 0.6; **Fig. 3A**). In the FuzDrop prediction profile, it was possible to identify three major uninterrupted peaks (*1*, *2*, and *3* in **Fig. 3A**) with very high P_DP_ scores (>0.8). Smaller LLPS-prone regions were identified at the ends of the CTD (*white squares*) and within the NTD (*black asterisks*). Such a distinctly bipartite distribution of low and high scores across the two domains was also clearly visible when per-residue P_DP_ scores were visualized in pseudo-colors on the AlphaFold structural model of CDKL5 (**Fig. 3B**).

**Figure 3.**
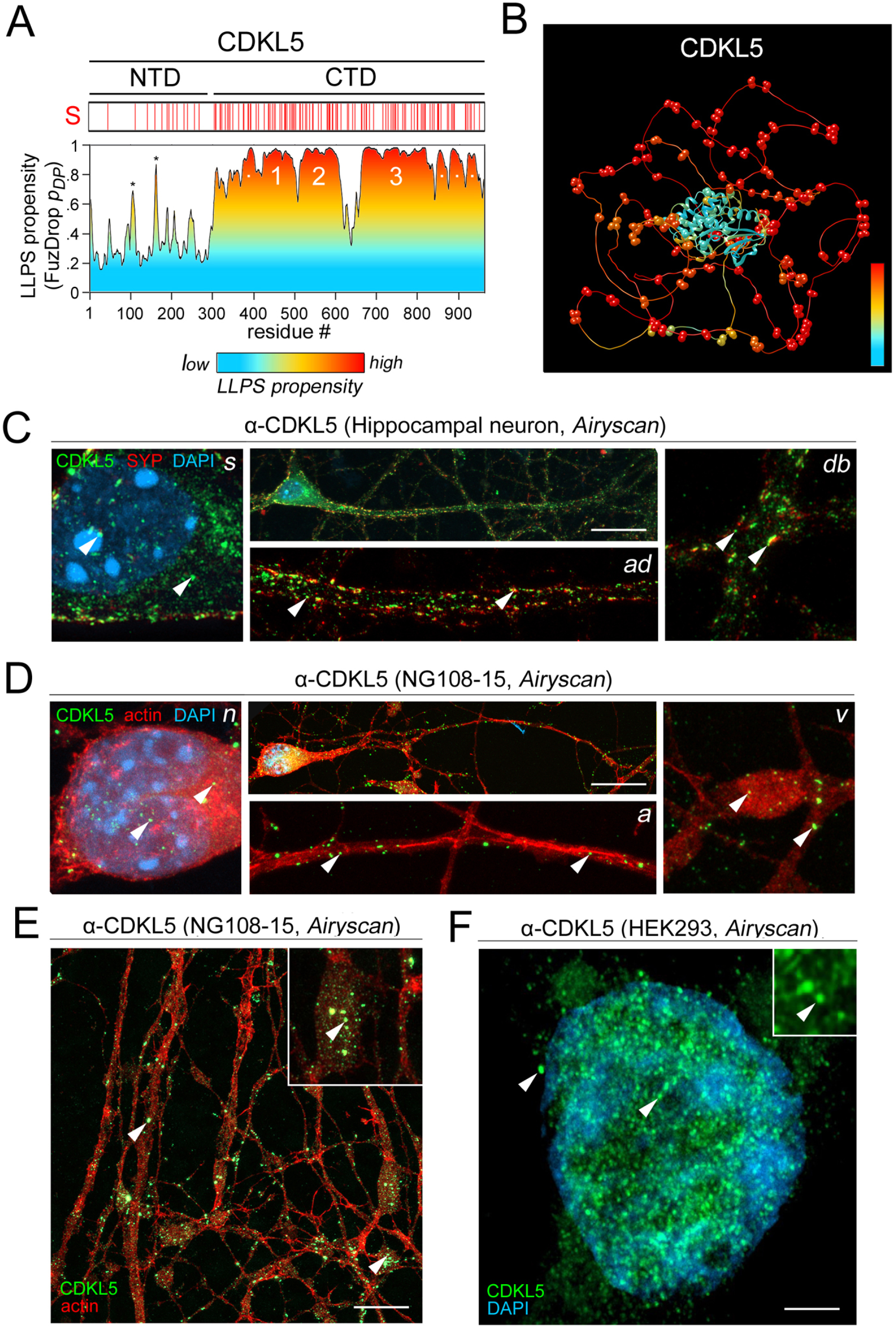
Endogenous CDKL5 forms condensate-like foci in neuronal and non-neuronal cells. **A**. The *lower graph* plots the per-residue FuzDrop LLPS propensity score (P_DP_ score) along the CDKL5 primary sequence. The area under the curve is colored using a *blue to red gradient* to highlight the protein regions with the highest LLPS propensity (P_DP_ score > 0.6; *orange*/*red*). The *white bar* above represents the protein with serine residues highlighted in *red*. The serine-rich CTD contains three extended regions with high LLPS propensity (1-3) and minor peaks marked by *white dots*. A few high-propensity peaks are also present within the NTD (*asterisks*). **B.** AlphaFold structural model of human CDKL5, as in Fig. 1C, pseudo-colored to display the FuzDrop per-residue LLPS propensity score using the same color scale as in *panel A*. The random coil serine-rich CTD, surrounding the central catalytic domain, is the region with the highest scores (*orange*/*red*). **C.** *Central upper panel*. Super-resolution (Airyscan, Zeiss) confocal fluorescence micrograph of a mouse primary hippocampal neuron immunostained to detect endogenous CDKL5 (*green*) and the presynaptic marker synaptophysin 1 (SYP, *red*). The nucleus was stained using DAPI (*blue*). The *left*, *lower middle,* and *right panels* show, respectively, magnified details of the neuronal somatic compartment (*s*), the proximal part of the apical dendrite (*ad*), and a more distal dendritic branch (*db*). Both intra- and extra-nuclear condensate-like foci (*arrowheads*) are visible. Scale bar: 20 μm. **D.** As in *C* for a differentiated NG108-15 neuronal cell. The *red* staining, obtained using fluorescently labelled phalloidin, marks here actin. The *middle lower panel* shows here a detail of the axon (*a*) and the *right panel* displays a neuritic varicosity (*v*). *Arrowheads* highlight condensate-like foci. Scale bar: 20 μm. **E.** Neuropile formed by intermeshed neurites, with varicosities, of differentiated NG108-15 cells. Note the enrichment of CDKL5 foci in varicosities. The *inset* in the *upper right corner* shows a magnified detail of the image encompassing a varicosity. The *arrowhead* indicates one of the condensate-like foci. Scale bar: 10 μm. **F.** CDKL5 immunostaining of a HEK293 cell (*green*). DAPI (*blue*) marks the nucleus. *Arrowheads* indicate condensate-like foci. Note the higher concentration of condensates in the nucleus compared to the cytoplasm. The *inset* in the *upper right corner* shows a magnified detail from a single confocal plane. Scale bar: 10 μm.

A similar analysis was performed for the other members of the CDKL family. CDKL-1, -2, and -4 all displayed low pLLPS scores, consistent with their shorter CTDs (**Suppl. Fig. 3A-B**). Only CDKL3, which has an intermediate length CTD, displayed an overall borderline score of 0.59, still slightly below the software prediction threshold. Notably, these scores correlated with the density of serine residues in the CTDs of the five kinases (r = 0.95, n = 5, p<0.02; **Suppl. Fig. 3B**). However, some regions of the CTD of CDKL2, CDKL4, and especially CDKL3, displayed above-threshold per-residue P_DP_ scores suggesting the possibility that they may have some degree of LLPS propensity. Interestingly, also the N/S-rich and A/Q-rich tails of the CDKL kinases in *C. elegans* and *D. melanogaster*, respectively, display above-threshold P_DP_ scores (**Suppl. Fig. 2A**). This observation further supports the notion that compositionally distinct, but similarly LLPS-prone, LCRs may have evolved in a convergent manner across clades (Vaglietti et al., 2023).

Taken together, these findings identify the long serine-rich LCR of CDKL5 as a strong candidate mediator of LLPS. The AlphaFold structural model also suggests that this CDKL5 region may form an ‘allosteric shell’ (Taylor et al., 2022) regulating the catalytic activity of the protein and mediating its homo- and hetero-typic protein-protein interactions, including those driving LLPS.

### CDKL5 forms intracellular LLPS-driven condensates in neuronal and non-neuronal cells

To determine whether CDKL5 undergoes LLPS in the cellular context, we tested in different cell types whether the protein forms rounded, membraneless foci (‘condensates’), as typical of LLPS-prone proteins, and whether these foci display distinctive properties of LLPS-driven protein assemblies.

By means of immunocytochemistry and Airyscan super-resolution confocal microscopy, we analyzed the subcellular distribution of CDKL5 in cultured neuronal (primary hippocampal neurons and differentiated NG108-15 cells) and non-neuronal (HEK293) cells (**Fig. 3C-F**). We found that endogenous CDKL5 forms rounded foci both in the nucleus and cytoplasm of all these cell types. In neuronal cells, the foci were also present in the axonal and dendritic compartments. The relative proportion of intra- *vs*. extra-nuclear condensates varied with the cell type. In primary neurons (**Fig. 3C**) and differentiated NG108-15 cells (**Fig. 3D-E**), condensates were prevalently distributed in the cytoplasm, the axon and the dendritic tree. In HEK293 cells, they were more concentrated in nuclei, although they were also present in the cytoplasm (**Fig. 3F**). These findings are consistent with previous observations of cell type-specific subcellular distribution and nucleo-cytoplasmic shuttling of the protein (Lin et al., 2005: Rusconi et al., 2008; Oi et al., 2017). In primary hippocampal neurons, the dendritic CDKL5 condensates co-localized or were in close juxtaposition with clusters of the presynaptic marker protein synaptophysin, in agreement with the fact that CDKL5 is present in both pre- and post-synaptic compartments (**Fig. 3C**). Presynaptic varicosities formed by NG108-15 cells (Han et al., 1991; **Fig. 3D**, *right panel,* and **Fig. 3E**) were particularly enriched in CDKL5 condensates.

Exogenously expressed, GFP-tagged CDKL5 formed similarly rounded foci in cells, displaying distribution patterns that matched those of the endogenous protein (**Fig. 4A-C**). These foci increased in size, also by coalescence, in a time- and expression level-dependent manner, as generally observed for LLPS-driven condensates (Rebane et al., 2020; *see also below*). Such coalescence was more pronounced in HEK293 cells (**Fig. 4C**) than in differentiated neuronal NG108-15 cells (**Fig. 4A**). In these neuronal cells, the condensates remained generally smaller and more distributed through the neuritic tree, matching the distribution pattern of endogenous CDKL5. In either cell type, both endogenous and exogenously expressed CDKL5 displayed variable distribution patterns and degrees of condensation across cells, likely related to cell cycle phases in mitotic HEK293 cells (Barbiero et al., 2017) or differentiation stages in neuronal NG108-15 cells.

**Figure 4.**
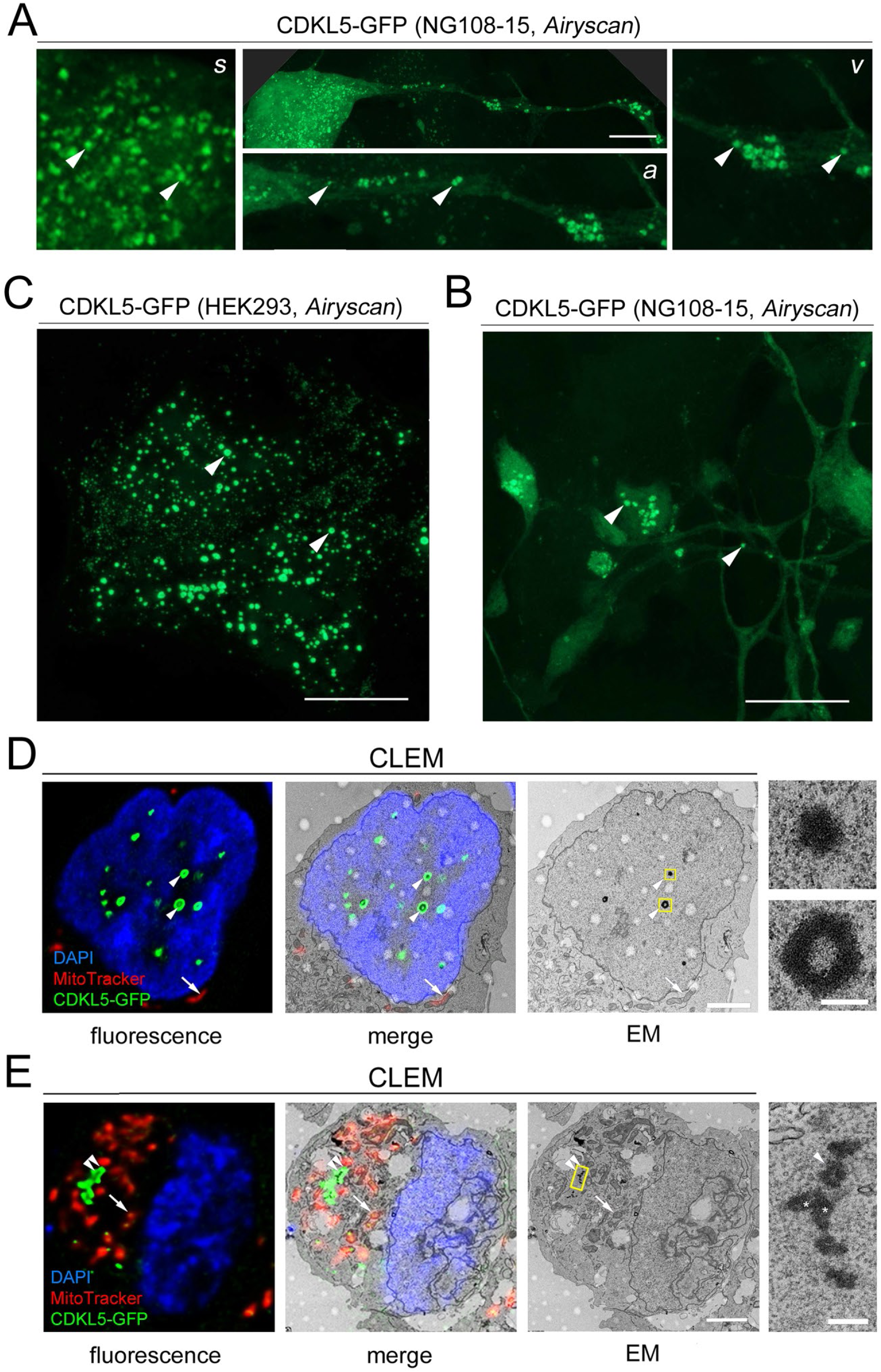
Exogenous CDKL5-GFP forms condensate-like foci in neuronal and non-neuronal cells. **A.** *Central upper panel*. Super-resolution (Airyscan, Zeiss) confocal fluorescence micrograph of a differentiated NG108-15 neuronal cell expressing CDKL5-GFP. The *left*, *lower middle,* and *right panels* show, respectively, magnified details of the somatic compartment (*s*), the axon (a), and a varicosity (*v*). Note the intra- and extra-nuclear condensate-like foci (*arrowheads*). Scale bar: 10 μm. The image upper corners have been filled in dark gray for graphical purposes. **B.** Neuropile of intermeshed neurites, with varicosities, of NG108-15 cells expressing CDKL5-GFP. *Arrowheads* indicate condensate-like CDKL5 foci in a varicosity (*left*) and along a neurite (*right*). Scale bar: 10 μm. **C.** Two HEK293 cells expressing CDKL5-GFP. *Arrowheads* indicate condensates. Scale bar: 10 μm. **D.** Correlative light and electron microscopy (CLEM) images of a HEK293 cell expressing CDKL5-GFP. The *first panel from the left* shows a confocal fluorescence image with intranuclear CDKL5-GFP condensates (*green*). The nucleus is stained with DAPI (*blue*) and mitochondria (*white arrow*) are marked by the MitoTracker dye (*red*). The *third panel* shows an EM image of a cell section at the same level of the confocal fluorescence image. The *second panel* is an overlay of the fluorescence and EM images. The two fluorescent condensate-like foci are highlighted by *arrowheads* and correspond to denser areas (boxed in *yellow* in the EM image) within the nuclear compartment, which are magnified in the two *right panels*. The MitoTracker signal overlaps with mitochondria, as expected. The two condensates are membraneless, and the larger one has an internal cavity also visible in the fluorescence image (see *Results*). Scale bars: 2.5 μm (larger images), 250 nm (smaller panels). **E**. As in *panel D*, for cytoplasmic condensate-like foci. The *fourth image on the right* illustrates a cluster of them (see *yellow box* in the EM image), two of which (*asterisks*) appear to be coalescing with each other. Scale bars as in *panel D*.

These findings provide morphological evidence that both endogenous and exogenously expressed CDKL5 form discrete, rounded intracellular foci as typical of proteins undergoing LLPS (Shin et al., 2017; Vaglietti et al., 2023).

To assess whether the CDKL5 intracellular foci are membraneless structures, as characteristic of LLPS-driven condensates, we performed *correlative light and electron microscopy* (CLEM) ultrastructural analyses of cells expressing CDKL5-GFP. CLEM scans, both in the nucleus (**Fig. 4D**) and in the cytoplasm (**Fig. 4E**), show that these structures are not delimited by membranes. Some of the foci were found in small clusters, and some of them appeared to be in the process of coalescing with each other (**Fig. 4E**, *right panel*), a hallmark feature of LLPS-driven condensates (Vaglietti et al., 2023). Moreover, some large foci appeared to be hollow (**Fig. 4D**, *right panel*), displaying an internal cavity, visible in both confocal fluorescence images and electron microscopy scans. Interestingly, morphologically similar structures have been recurrently observed in the condensates formed by LLPS-prone proteins, such as TDP-43 and Oskar. This configuration of the condensates drives from their internal partitioning related to local changes in the physico-chemical environment or to the presence of co-separated molecular components, such as other proteins and/or nucleic acids (Kistler et al., 2018; Schmidt and Rohatgi, 2016; Dai and Yang, 2024).

Finally, we tested whether, in addition to their morphology and ultrastructure, the intracellular foci formed by CDKL5 display additional hallmark features of LLPS-driven protein condensates (Sabari et al., 2018; Basu et al., 2020).

We first tested the dynamics of CDKL5 molecular diffusion between the condensates and the surrounding cellular environment by measuring *fluorescence recovery after photobleaching* (FRAP; **Fig. 5A)**. These experiments revealed rapid FRAP within 60 s, which could be fitted to an exponential curve with a *t*-half of 12.2 s and a mobile fraction of 74% (mean values of n = 11 condensates), indicative of rapid molecular exchange between the foci and the surrounding environment, in the range of other LLPS-prone proteins (e.g. Shin et al., 2017; Fan et al., 2020; Vaglietti et al., 2023).

**Figure 5.**
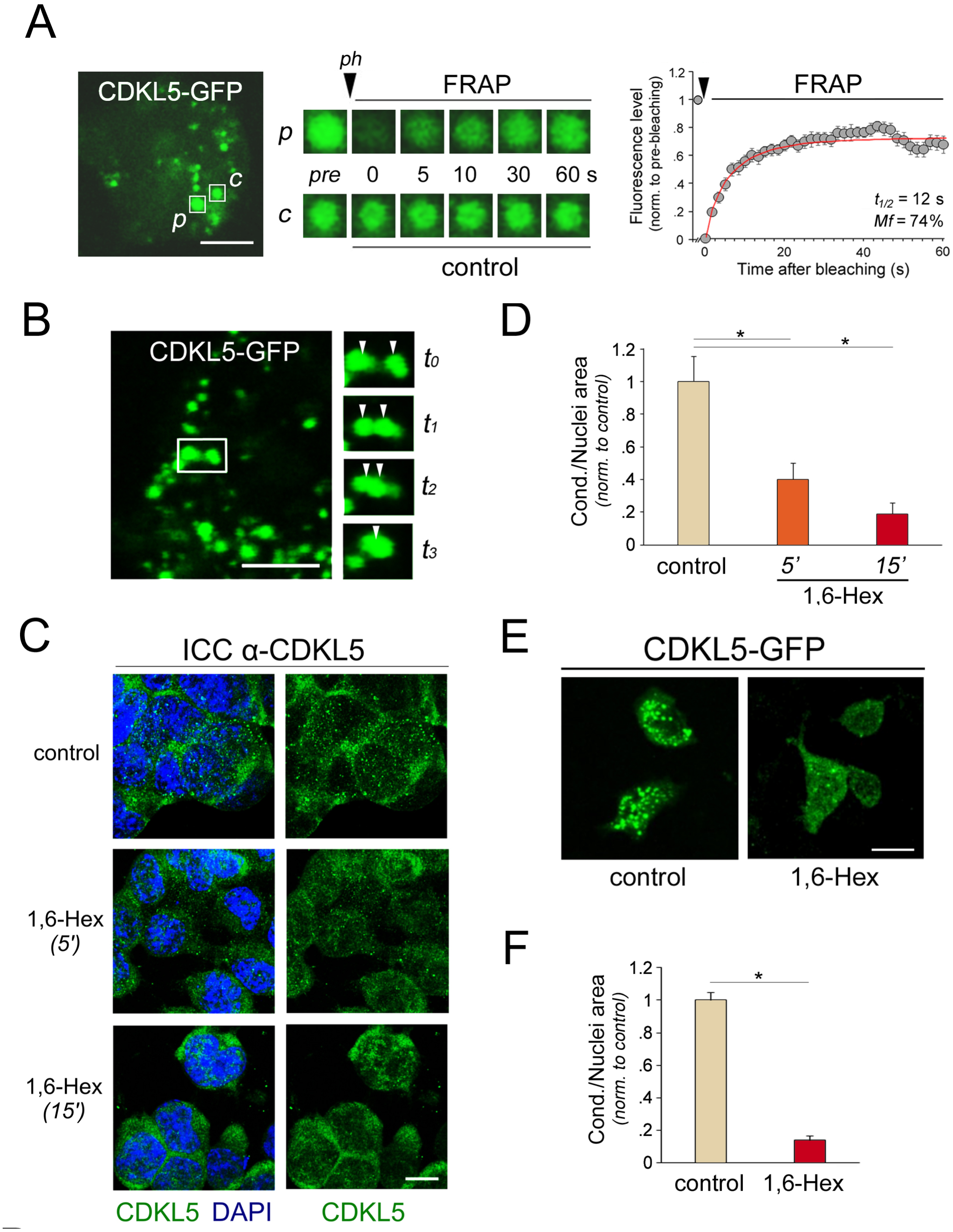
The CDKL5 foci display hallmark features of LLPS-driven condensates. **A.** FRAP experiments on CDKL5 foci. The *left panel* shows a confocal image of a HEK293 cell expressing CDKL5-GFP. Scale bar: 5 μm. Two condensates are highlighted within *white squares*. One of them was photobleached (*p*) and the other one was used as a control (*c*). In the *middle section*, the *upper series* of time-lapse images displays the condensate *p* before (*pre*) and at the indicated times after photobleaching. Condensate *p* substantially recovered its fluorescence in about a minute after photobleaching. For condensate *c* (*lower series*), which was not actively photobleached, only modest fluorescence bleaching was observed due to repeated imaging. The *right graph* displays the mean FRAP time course for photobleached CDKL5-GFP foci (n = 11; half-time (*t_1/2_*) = 12 s; mobile fraction (*Mf*) = 74%. **B.** The *left panel* displays a single confocal section of a HEK293 cell expressing CDKL5-GFP with two neighboring foci, highlighted within the *white box*. Scale bar: 4 μm. *On the right*, frames of a time-lapse live-cell imaging experiment showing, at four time points (*t_0_*-*t_3_*) spaced 2 seconds apart, how the two foci (*arrowheads*) rapidly coalesce into a larger one (*arrowhead*). **C.** Confocal fluorescence images of NG108-15 cells immunostained to detect the distribution of endogenous CDKL5 (*green*). The DAPI *blue* signal in the *left column* of images marks cell nuclei. The cells were fixed after 5 or 15 min of 1,6-Hexanendiol (*1,6-Hex*) treatment. Mock-treated cells were used as the *control* group. Note how 1,6-Hex rapidly promotes the dissolution of CDKL5 foci. Scale bar: 10 μm. **D.** Quantification of the effect of 1,6-Hex on endogenous CDKL5 condensate-like foci, as shown in *panel C*. The mean CDKL5-positive area occupied by these structures is significantly reduced by 1,6-Hex. Values are normalized to the mean of the control group. *Asterisks* indicate statistically significant differences. **E.** Confocal fluorescence images of HEK293 expressing CDKL5-GFP (*green*). The image *on the right* shows cells fixed after 15 min of 1,6-Hex treatment. Untreated *control* cells are shown *on the left*. Note how 1,6-Hex rapidly promotes the dissolution of condensate-like CDKL5-GFP foci. Scale bar: 10 μm. **F.** Quantification of the effect of 1,6-Hex, as shown in *panel E*. The mean proportion of the CDKL5-positive area occupied by condensate-like foci is significantly reduced by 1,6-HEX. Values are normalized to the mean of the control group. *Asterisks* indicate statistically significant differences.

Second, in live-cell imaging experiments, we observed that smaller CDKL5-GFP condensates tend to coalesce into larger ones in a time-dependent manner, another typical feature of LLPS-driven, liquid-like condensates (**Fig. 5B**).

Third, we tested whether the CDKL5 foci are sensitive to 1,6-Hexanediol (1,6-Hex; **Fig. 5C**), a compound known to disrupt the relatively weak intermolecular interactions underlying LLPS (Ulianov et al., 2021; Vaglietti et al., 2023). Towards this aim, we compared the condensation of endogenous CDKL5 in NG108-15 cells treated with either 1,6-Hex (for 5 or 15 min), or mock-treated with only the vehicle (DMEM), and then immunostained to detect CDKL5. We found that 1,6-Hex caused an overall significant reduction in the fraction of cell area occupied by condensates (F_(2,44)_=11.78, p<0.001, one-way ANOVA; **Fig. 5D**). Indeed, already at 5 min, the area occupied by condensates was reduced to 39.92 ± 9.86 % (n = 15 microscopy fields) of the same area measured under control condition (100 ± 15.45 %, n = 19, p<0.01, Newman-Keuls (NK) *post hoc* test; data normalized to the control group mean). This reduction was even more pronounced at 15 min (18.83 ± 6.59 %; n = 13, p<0.01 NK *post hoc* test *vs.* control group).

A similar sensitivity to 1,6-Hex was also observed for condensates formed by exogenously expressed CDKL5-GFP in HEK293 cells (**Fig. 5E**). Indeed, 1,6-Hex application for 15 min induced a ∽86% reduction of the area occupied by condensates in comparison with control conditions (100 ± 4 %, n = 64 fields, *vs* 13 ± 2 %, n = 47, respectively, p<0.001, *t*-test; **Fig. 5F**).

Taken together, these findings indicate that CDKL5 foci in the cellular context display morphological, ultrastructural, dynamic, and biochemical hallmark features of LLPS-driven condensates.

### The serine-rich CTD has a prominent role in mediating CDKL5 LLPS

To determine which portion of the CDKL5 protein may drive LLPS, we started by comparing the relative ability to undergo spontaneous condensation of GFP-tagged full-length (FL) CDKL5 (as a positive control), of its two major NTD and CTD fragments, and of GFP alone (as a negative control), in both neuronal and non-neuronal cells (**Fig. 6A,B**).

**Figure 6.**
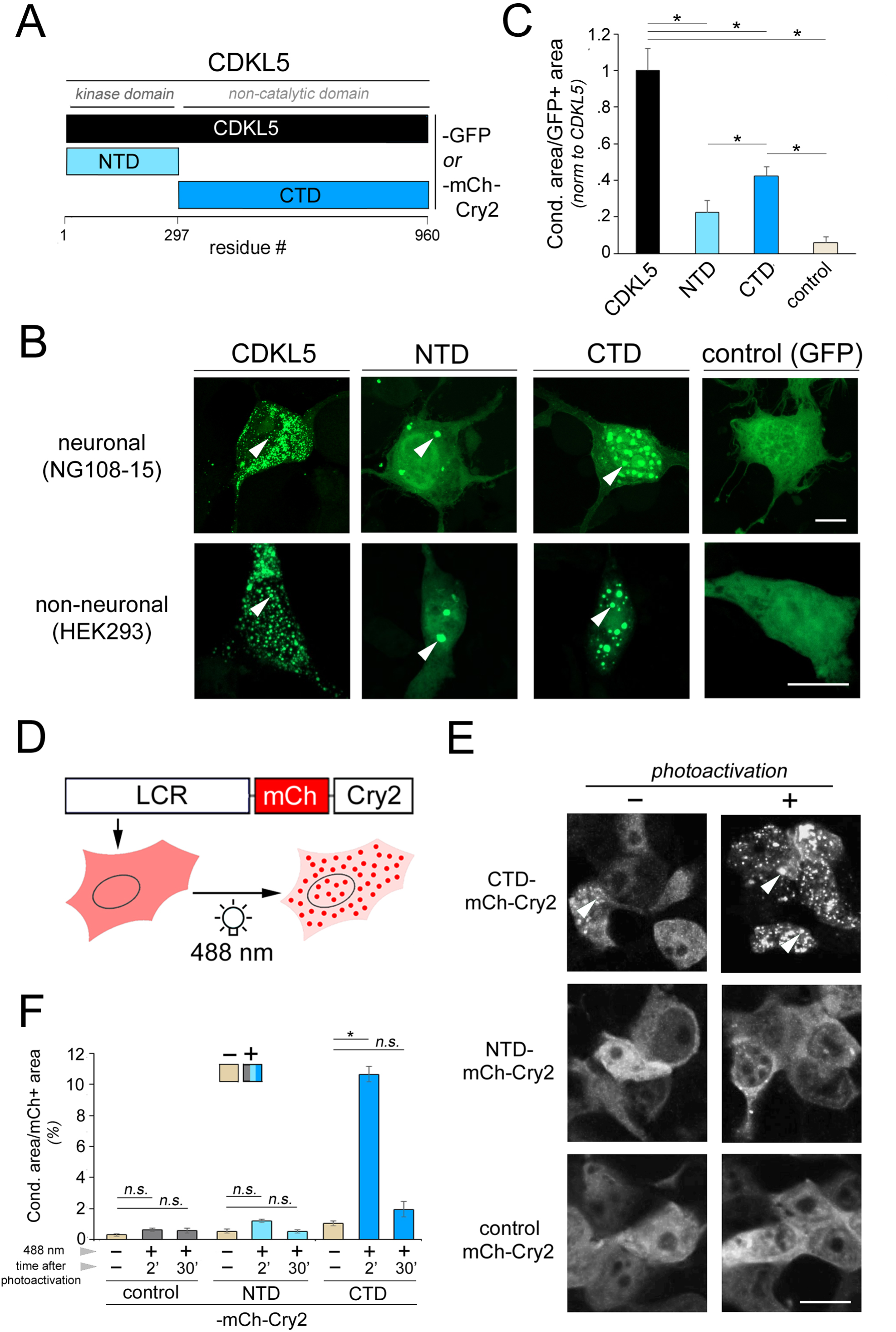
CDKL5 condensation is primarily driven by its CTD. **A.** Schematic representation of the constructs for the cellular expression of full-length CDKL5 (*black*), and of its NTD (*cyan*), or CTD (*turquoise*), as fusion proteins with either GFP or mCh-Cry2 for the study of spontaneous and triggered LLPS, respectively. **B.** GFP-tagged CDKL5, its NTD or CTD fragments, or GFP alone, were expressed in neuronal (differentiated NG108-15, *upper row*) and non-neuronal (HEK293, *lower row*) cell to study their spontaneous condensation (*arrowheads* indicate condensates). While CDKL5-GFP forms a myriad of small condensates, the NTD displays only a few larger condensates, whereas the CTD forms numerous condensates of intermediate size often concentrated in the nucleus, especially in NG108-15 cells. Scale bars: 10 μm. **C.** Quantification of the spontaneous condensation of the different constructs expressed in HEK293 cells as shown in *panel B*. Values are normalized to the mean of the CDKL5-GFP group. *Asterisks* indicate statistically significant differences. **D.** Schematic representation of the optoDroplet system for studying rapidly triggered LLPS (Shin et al., 2017). Cell cultures expressing an LLPS-prone peptide of interest, such as an LCR, in fusion with mCh-Cry2, and control cultures expressing mCh-Cry2 alone, are illuminated with blue light (488 nm) to optically trigger the rapid formation of condensates (‘optoDroplets’, i.e. the *red dots* in the right cell). Control cultures expressing mCh-Cry2 display negligible condensation upon photoactivation. mCh fluorescence is used to monitor the fusion protein localization. **E.** Confocal fluorescence images of cell cultures expressing CDKL5 NTD or CTD fused to mCh-Cry2, or mCh-Cry2 alone (*control*). For each expressed construct, some cultures were photoactivated by a pulse of 488 nm blue light (*right column*, ‘+’) while some other control cultures were not photoactivated (*left column*, ‘-’). *Arrowheads* indicate condensates. Scale bar: 10 μm. **F.** Bar graph showing the proportion (%) of the mCh-positive cell area occupied by condensates in cell cultures treated as shown in *panel E*. For each indicated construct, condensation was quantified in control, non-photoactivated cultures (basal spontaneous condensation), and in cultures that were photoactivated (triggered condensation) and fixed at either 2 or 30 min after photoactivation. *Asterisks and ‘n.s.’* indicate statistically significant and non-significant differences, respectively, between experimental groups of interest. Note that only the CTD construct undergoes a significant induction of condensation at 2 minutes after photoactivation. Such condensation is reversible and essentially disappears within 30 min. Scale bar: 5 μm.

When we compared the cell area occupied by condensates in transfected cells of the four experimental groups, we observed significant overall differences in a one-way ANOVA statistical analysis (F_(3,105)_ = 19.617, p < 0.001; **Fig. 6C**). Pairwise *post hoc* comparisons between the groups revealed that the CTD fragment displayed significant condensation compared to the GFP negative control group (42.45 ± 4.86%, n = 31 microscopy fields, *versus* 5.80 ± 3.42%, n = 11, respectively; values normalized to the mean of the FL positive control group; p < 0.04, NK test), similar to what observed for the FL protein (100 ± 12.25%, n = 35; p < 0.001 *versus* the GFP group, NK test). Conversely, the NTD group, while displaying noticeable formation of large condensates in some cells (relative condensate area in cells 22.72 ± 6.15%, n = 32, values normalized to the FL group), did not show a significant difference from the GFP group (p = 0.24). Both the CTD and the NTD displayed significantly lower condensation rates compared to the FL protein (p < 0.001 in both cases).

We also tested the relative ability of the NTD and the CTD fragments to undergo rapid, triggered LLPS using the optoDroplet system (Shin et al., 2017), an optogenetic tool allowing to phototrigger the condensation of LLPS-prone proteins/peptides in a spatially and temporally controlled manner in the cellular environment (**Fig. 6D**). In these experiments, the NTD or the CTD were expressed in cells in fusion with the red-fluorescent protein mCherry, for protein visualization, and an *A. thaliana* Cry2 fragment (mCh-Cry2) to confer photoactivability (Shin et al., 2017). mCh-Cry2 alone was expressed in control cultures (**Fig. 6E**).

Cell cultures expressing NTD-mCh-Cry2, CTD-mCh-Cry2, or mCh-Cry2, were either photoactivated using 488 nm blue light for 2 min (Shin et al., 2017; see *Methods*) or left untreated as a control, and then fixed at 2 min (to test for LLPS induction) or 30 min (to test for rapid LLPS reversibility) after the end of photoactivation. For each construct, we quantified the relative proportion (%) of mCh-positive cell area occupied by condensates in cultures that were either photoactivated (‘*+’*) or not (‘-*’*, *control*; **Fig. 6F**). A two-way ANOVA revealed overall significant effects of the construct (F_(2,414)_=123.61, p < 0.001), of photoactivation (F_(2,414)_=90.81, p < 0.001), and their interaction (F_(4,414)_=68.83, p < 0.001; n = 44-48 microscopy fields per group). *Post hoc* comparisons showed that the mCh-Cry2 control construct alone did not undergo significant condensation after photoactivation, as expected (p = 0.96, NK *post hoc* test). Conversely, the CTD-mCh-Cry2 construct displayed, even in basal conditions, some degree of spontaneous condensation which was strongly increased upon photoactivation (p < 0.001, NK *post hoc* test). The NTD-mCh-Cry2 construct underwent instead only a minimal, non-significant increase in condensation upon photoactivation (p = 0.67, NK *post hoc* test). The phototriggered condensation of the CTD was rapidly reversible, as typical of LLPS-driven processes (Vaglietti et al., 2023). Indeed, the condensation state of the CTD-mCh-Cry2 construct was not different from that of control cultures 30 minutes after photoactivation (p = 0.17, NK *post hoc* test).

Taken together, these analyses indicate that the physiological condensation pattern of the full-length protein depends on the presence of both the NTD and the CTD, and that the latter appears to be primarily responsible for driving CDKL5 LLPS.

### A defined serine-rich CTD internal fragment (CTIF) has a pivotal role in driving LLPS

The CTD constitutes ∼2/3 of the entire CDKL5 protein, for a total of 663 residues in the isoform most expressed in the brain, encompassing residues 298-960. The previous findings prompted us to test whether the entire CTD is necessary to mediate LLPS or a specific subregion of it may play a more prominent role in driving this process.

In search of candidate LLPS-mediating CTD subregions, we considered that previous reports based on CDKL5 deletions indicated that residues between 525 and 781 have an important role in regulating the subcellular distribution of the protein (Ricciardi et al., 2009), which, from the perspective of our analysis, may also have to do with LLPS. Remarkably, we found that this region of the protein has substantial overlap with the longest uninterrupted stretch of residues, between 657 and 840, with very high LLPS propensity in the FuzDrop prediction diagram (P_DP_ score > 0.8; peak *3* in **Fig. 7A**). Indeed, the average per-residue P_DP_ score in this region (0.93 ± 0.01, n = 184 residues), was significantly higher than the average score across residues of the upstream (‘CTDu’; a.a. 298-656; 0.83 ± 0.01, n = 395 residues) and downstream (‘CTDd’; a.a. 841-960; 0.86 ± 0.01, n=120 residues) parts of the CTD, and of the NTD (0.33 ± 0.01, n=297). A one-way ANOVA revealed a significant overall difference among the four protein segments (F_(3,956)_ = 1162.4, p < 0.001). Furthermore, pairwise *post hoc* tests highlighted significant differences between the 657-840 fragment and each one of the three other fragments, i.e. CTDu, CTDd, and NTD (p < 0.001, NK test, in all instances; **Fig. 7B**).

**Figure 7.**
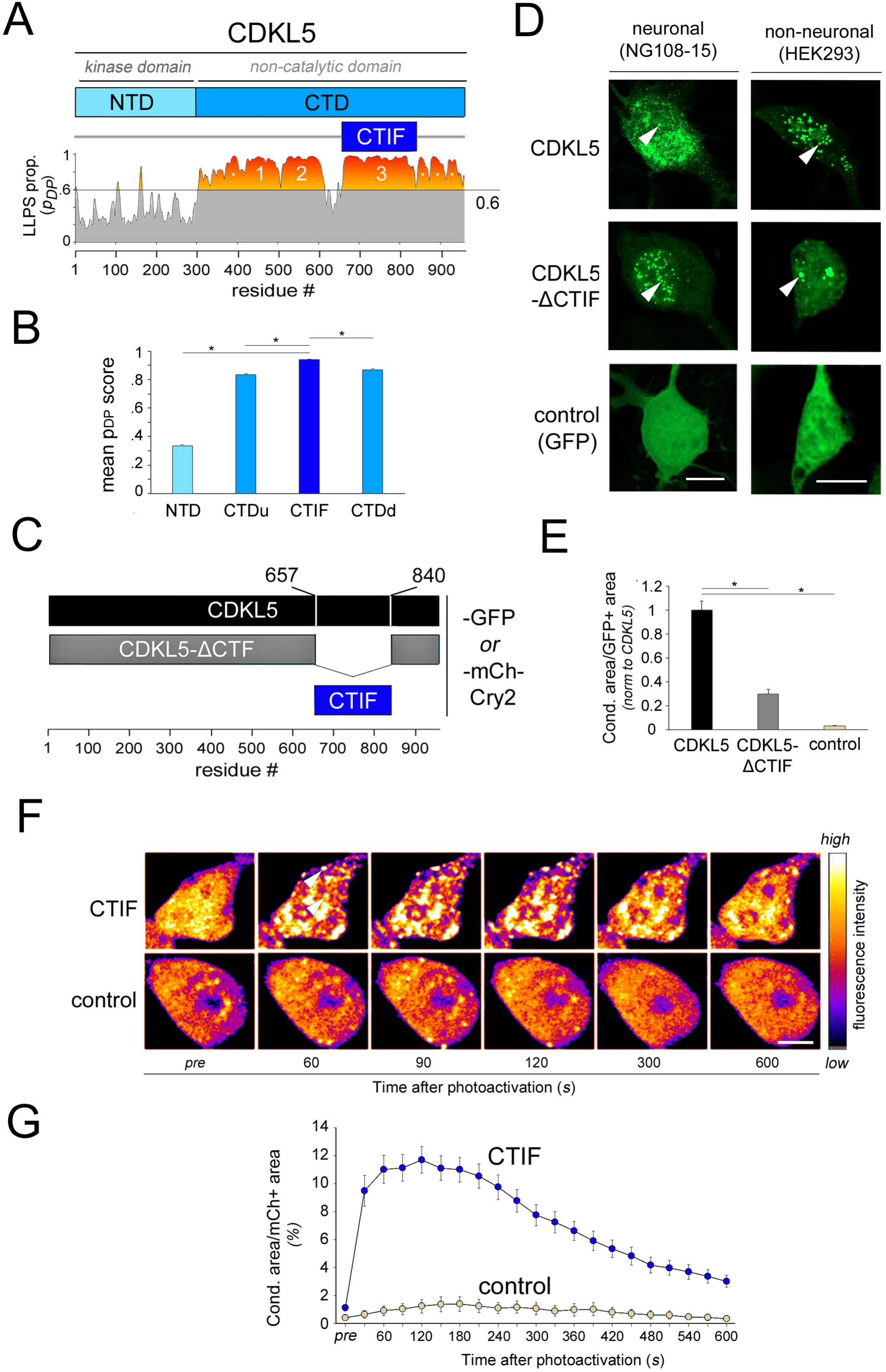
A C-terminal internal fragment (CTIF) of the CTD has a key role in driving CDKL5 LLPS. **A**. The *upper bar* displays a schematic representation of the CDKL5 primary sequence. The NTD is highlighted in *cyan* and the CTD in *turquoise*. The *lower graph* plots the FuzDrop per-residues LLPS propensity score (P_DP_ score) along the primary sequence of human CDKL5. The area under the curve with the above-threshold LLPS propensity (P_DP_ score > 0.6) is highlighted in *orange*/*red*. The *blue bar* tract identifies a C-terminal internal fragment (CTIF; a.a. 657-840) of CDKL5 corresponding to the longest uninterrupted region of the CTD with very high (>0.8) P_DP_ score, (i.e. peak 3, see Fig. 3A). **B.** Bar graph plotting the mean P_DP_ score across residues of the NTD (a.a. 1-297), the CTIF (a.a. 657-840), and the two upstream (*u*) and downstream (*d*) CTD portions, i.e. CTDu (a.a. 298-656) and CTDd (a.a. 841-960), respectively. Note how the CTIF has the highest mean LLPS propensity score. **C.** Schematic representation of the constructs that were generated for the cellular expression of full-length CDKL5 (*black*), or a CTIF deletion mutant (CDKL5-ΔCTIF, *gray*), or the CTIF alone (*blue*), as fusion proteins with either GFP or mCh-Cry2 for the studyS of spontaneous and triggered LLPS, respectively. **D.** GFP-tagged CDKL5, its CDKL5-ΔCTIF deletion mutant, or GFP alone as a control, were expressed in neuronal (differentiated NG108-15 cells, *left columna row*) and non-neuronal (HEK293 cells, *right column*) to study their spontaneous tendency to form condensates (*arrowheads*). CDKL5-ΔCTIF can still form condensates in both cell types, although at a significantly lower level than the full-length CDKL5 protein. Scale bars: 10 μm. **E.** Quantification of the spontaneous condensation of the different constructs expressed in HEK293 cells, as shown in *panel D*, as the mean proportion of the GFP-positive cell area occupied by condensates. Values are normalized to the mean of the CDKL5-GFP group. *Asterisks* indicate significant differences. CTIF deletion significantly reduces condensation. **F**. Confocal fluorescence images of cell cultures expressing either CTIF-mCh-Cry2 or mCh-Cry2 alone (*control*) at the indicated time points before (*pre*) and after photoactivation in live-cell imaging experiments. Photoactivation induces massive condensation of CTIF-mCh-Cry2 (but not of mCh-Cry2 alone) which peaks at 2 min after the end of the 488 nm light pulse and starts declining at later time points. *Arrowheads* indicate condensates. Fluorescence intensity represented using the pseudocolor scale shown *on the right*. **G.** Graph plotting the mean proportion of mCh-positive cell area occupied by condensates in cells expressing CTIF-mCh-Cry2 or mCh-Cry2 at the indicated time points of live-cell optoDroplet induction experiments, as shown in *panel F*.

Thus, we hypothesized that this CTD internal fragment (‘CTIF’) may play a pivotal role in driving LLPS. To test this hypothesis, we expressed, in both neuronal and non-neuronal cells, a GFP-tagged form of CDKL5 devoid of the CTF (CDKL5-ΔCTF; **Fig. 7C**) and assessed whether such an internal deletion may impact the ability of the protein to form intracellular condensates. We found that CDKL5-ΔCTF was still able to form condensates, but to a significantly lower extent than wild type CDKL5 in both neuronal and non-neuronal cells (**Fig. 7D**). In both cell types, the protein appeared to condensate more frequently in the nuclear compartment.

A one-way ANOVA indicated overall significant differences between the experimental groups in HEK293 cells (F_(2,72)_=39.751, p < 0.001; **Fig. 7E**). The CDKL5-ΔCTF condensates occupied a significantly lower proportion of the cellular area than the condensates formed by wild-type CDKL5 (29.6 ± 3.88 %, n = 48 microscopy fields, *vs.* 100 ± 7.17 %, n = 44; p < 0.01, NK *post hoc* test; values normalized to the wild-type CDKL5 group).

To further confirm the LLPS propensity of the CTIF alone, we studied its ability to undergo rapid phototriggered phase separation when expressed in HEK293 cells as a CTIF-mCh-Cry2 fusion protein (**Fig. 7F**). In live-cell experiments, we found that the CTIF was able to undergo robust LLPS upon photoactivation, which was not observed under the same conditions in control cells expressing mCh-Cry2 (**Fig. 7G**). A one-way ANOVA for repeated measures indicated significant overall differences between the two experimental groups in terms of construct, time, and their interaction (F_(20,1020)_ = 59.38, p < 0.001). *Post hoc* tests indicated that photoactivation induced significant condensation of mCh-Cry2-tagged CTIF in comparison with basal conditions (peaking at 2 min after photoactivation; p < 0.001 NK test) but not of mCh-Cry2 alone (p = 0.81). Remarkably, the CTIF-mCh-Cry2 condensation was reversible, as typical of LLPS-driven processes (e.g. Vaglietti et al., 2023), rapidly decreasing towards baseline levels within 10-15 min.

Taken together, these findings indicate that the serine-rich CTIF represents a key player in the phase transitions of CDKL5. As the CDKL5-ΔCTF was still able to condensate at reduced levels, the other parts of the CTD and/or the NTD must also play some role in CDKL5 LLPS.

### The serine-rich CTD is affected almost exclusively by disease-related truncating mutations

Given the largely divergent compositional and functional features of the CTD and the NTD, we analyzed quantitatively whether the type and frequency of disease-related coding mutations may be different in the gene exons encoding the two portions of the CDKL5 protein.

Towards this aim, we first derived a list of non-truncating (‘NTMs’, i.e. missense and in-frame indel) and truncating (‘TMs’, i.e. nonsense and frameshift) non-synonymous coding mutations, categorized as ‘pathogenic’, from the *ClinVar* database (**Suppl. Table 2; Fig. 8A**). Then, we calculated the relative proportion of the two mutation classes that affect the two CDKL5 domains (**Suppl Fig. 4**). We found that the CTD is essentially affected almost entirely by truncating mutations, whereas the NTD is targeted by both types of mutations (**Fig. 8A**). The differential distribution of TMs and NTMs across the NTD and the CTD is statistically significant (p < 0.01, Fisher’s exact test; **Suppl Fig. 4**).

**Figure 8.**
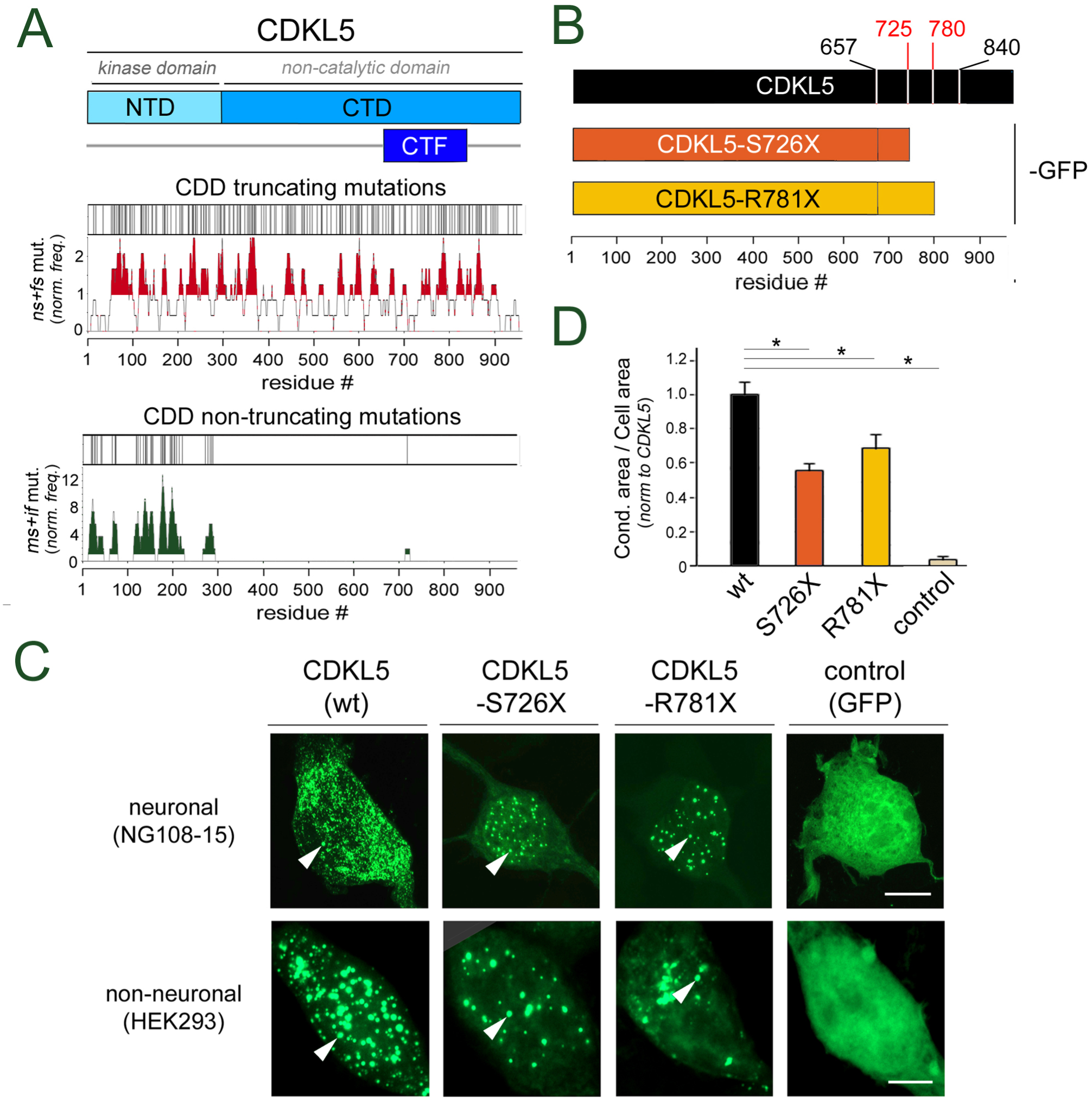
Disease-related mutations truncating the CDKL5 protein within the CTIF impair LLPS. **A**. *Upper two bars*. Schematic representation of the CDKL5 protein with the NTD (*cyan*), the CTD (*turquoise*), and the CTIF (*blue*). The *thin vertical lines* on the two white bars represent, as indicated, the positions of truncating (nonsense (*ns*) and frameshift (*fs*)) or non-truncating (missense (*ms*) and in-frame indel (*if*)) disease-related coding mutations along the CDKL5 open reading frame (ORF). The *graphs below* the white bars plot the local frequency of truncating and non-truncating disease-related mutations, as calculated in a sliding window of 21 triplets centered around each codon of the CDKL5 ORF. Values are normalized to the mean occurrence of the two types of mutations along the entire ORF. Above-average (>1) frequency peaks are highlighted in *red* and *dark green*, respectively, for truncating and non-truncating mutations. **B.** Schematic representation of the constructs that were generated for the cellular expression of wild-type CDKL5 (*black*) and two of its disease-related mutants (S726X and R781X) as fusion proteins with GFP for the study of spontaneous LLPS. **C.** GFP-tagged CDKL5, its two disease-related mutants, or GFP alone, were expressed in neuronal (NG108-15 cells, *upper row*) and non-neuronal (HEK293 cells, *lower row*) to study their spontaneous tendency to form condensates (*arrowheads*). Scale bar NG108-15: 10 μm. Scale bar HEK293: 5 μm). **D.** Quantification of the spontaneous condensation of the indicated constructs in HEK293 cells, as shown in *panel C*, expressed as the mean proportion of the GFP-positive cell area occupied by condensates. Values are normalized to the mean of the CDKL5-GFP group. *Asterisks* indicate statistically significant differences.

The fact that the pathogenic TMs can affect the CTD for almost its entire length strongly suggests that even the most distal portions of this domain are relevant to CDKL5 function.

### Disease-related truncations eliding the CTF and the downstream portion of the CTD impair CDKL5 LLPS and catalytic activity

To test whether the disease-related truncating mutations of the CTD may alter CDKL5 LLPS, we generated constructs for the expression of two GFP-tagged pathogenic mutants of CDKL5 in which the protein is truncated within the CTIF (i.e. S726X and R781X; Bertani et al., 2006, Rusconi et al., 2008; Trump et al., 2016; **Fig. 8B**). CDKL5-S726X and CDKL5-R781X lack, respectively, ∼2/3 and ∼1/3 of the CTIF together with the more distal portion of the CTD (a.a. 781-960).

Then, we compared the relative ability to undergo LLPS of wild-type CDKL5 (*CDKL5 wt*), its two truncated forms *CDKL5-S726X* and *CDKL5-R781X*, and GFP as a negative control (**Fig. 8C**). We found that both deletion mutants displayed a significant reduction in spontaneous condensation in comparison with full-length CDKL5 (**Fig. 8D**). Indeed, a one-way ANOVA showed an overall significant difference between the experimental groups (F_(3,225)_ = 30.644, p < 0.001). *Post hoc* tests indicated that the cell area occupied by condensates was significantly lower for both CDKL5-S726X (54.89 ± 4.34 %, n = 92 microscopy fields; p < 0.001, NK test) and CDKL5-R781X (69.11 ± 6.19 %, n = 34; p < 0.01, NK test) in comparison with full-length CDKL5 (100 ± 5.39 %, n = 34 microscopy fields; values normalized to the mean of the wt CDKL5 group). Both mutants still displayed significant condensation in comparison with the negative control group (GFP; p < 0.001, NK test, in both instances). The CDKL5-S726X mutant underwent a ∼14% greater reduction in condensation with respect to the other mutant CDKL5-R781X. Although this difference is not significant (p = 0.16, NK test), it suggests that a larger CTD deletion may cause a more pronounced LLPS impairment.

These observations indicate that the more or less complete loss of the CTIF (a.a. 657-840), together with the distal C-terminal portion of the CTD (a.a. 841-960), significantly impairs CDKL5 LLPS.

Consistent with this view, we found that the C-terminal fragments, i.e. CDKL5_726-960_ and CDKL5_781-960_ (**Fig. 9A**), which are elided by the two disease-related mutations, were both able to undergo condensation in the optoDroplet assay (**Fig. 9B**). A two-way ANOVA (**Fig. 9C**) indicated overall significant differences between the experimental groups in terms of construct (F_(2,416)_ = 77.44, p < 0.001), treatment (photoactivation; F_(2, 416)_ = 272.45, p < 0.001), and their interaction (F_(4,416)_ = 85.28, p < 0.001). *Post hoc* tests indicated that photoactivation was able to induce significant condensation of both the mCh-Cry2-tagged CDKL5_726-960_ and the CDKL5_781-960_ fragments (p < 0.001 in both instances *vs.* the respective non-photoactivated control groups, NK test) but not of mCh-Cry2 alone (control group, p = 0.69) at 2 min after photoactivation. Notably, the degree of condensation induced by photoactivation was significantly greater for the longer CTD terminal fragment (CDKL5_726-960_) than for the shorter one (CDKL5_781-960_; p < 0.001 NK test). As expected, the condensation of both constructs was rapidly reversible and reached baseline values within 30 min after photoactivation. Indeed, the condensation level at 30 minutes was not significantly different than in non-photoactivated control cultures for both constructs (p = 0.72 and p = 0.21, respectively, NK *post hoc* test).

**Figure 9.**
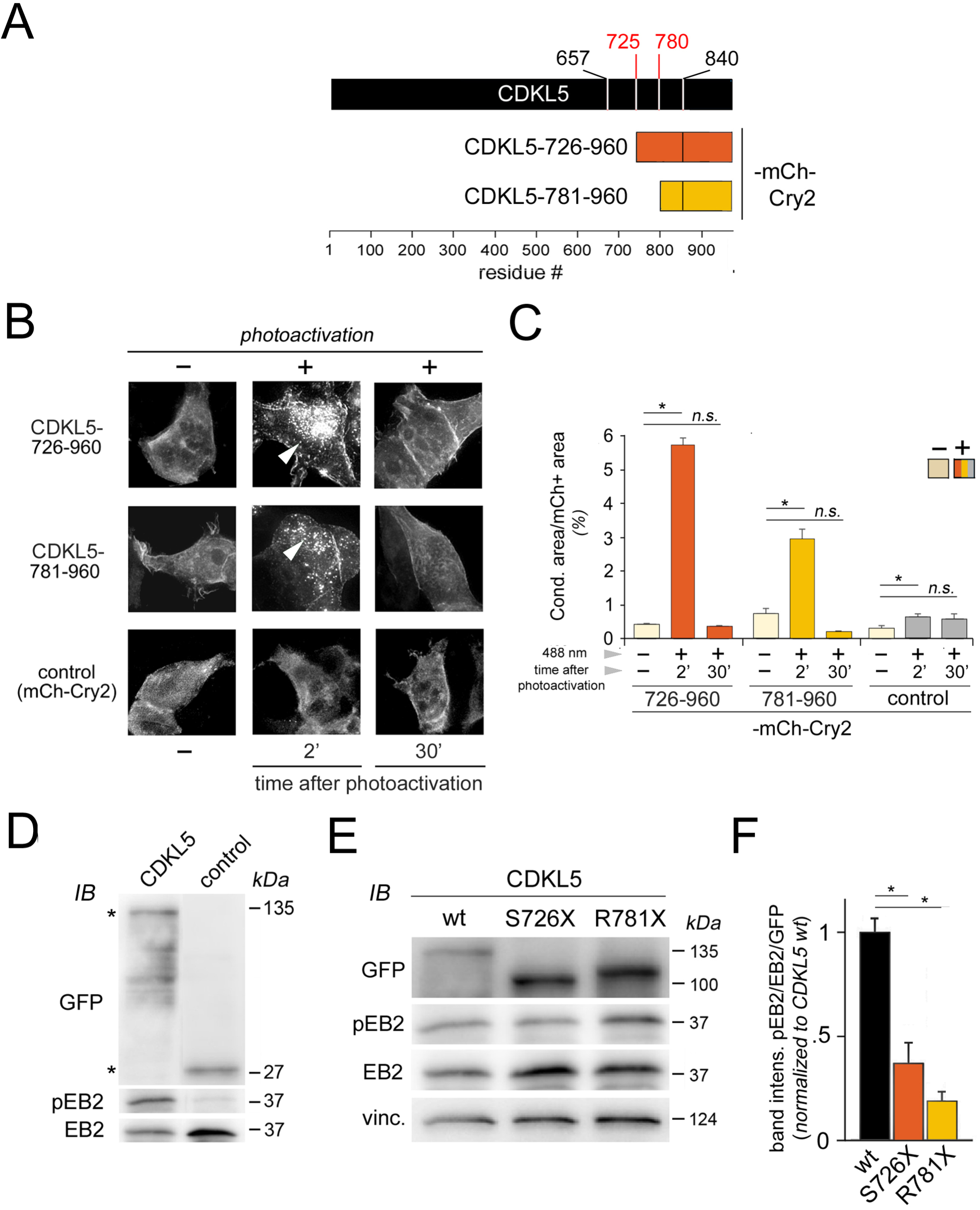
The C-terminal tails elided by truncating mutations can undergo LLPS and their loss impairs CDKL5 catalytic activity. **A.** Schematic representation of the constructs that were generated for the cellular expression of the C-terminal tails of CDKL5 (a.a. 726-960 or a.a. 781-960) as fusion proteins with mCh-Cry2 to study their LLPS propensity using the optoDroplet system. **B.** Confocal fluorescence images of cell cultures expressing the indicated mCh-Cry fusion proteins or mCh-Cry2 alone (*control*). For each expressed construct, some cultures were photoactivated with 488 nm blue light (*right column*, ‘+’) while control cultures were not photoactivated (*left column*, ‘-’). Photoactivated cultures (+) were fixed at either 2 or 30 min after photoactivation to assess, respectively, LLPS induction and reversibility. *Arrowheads* indicate condensates. Scale bar: 5 μm. **C.** Bar graph showing the proportion of the mCh-positive cell area occupied by condensates in cultures treated as shown in *panel B*. For each indicated construct, condensation was quantified in control, non-photoactivated cultures (basal spontaneous condensation), and in cultures that were photoactivated (triggered condensation) and fixed at 2 or 30 min after photoactivation. Statistically significant (*asterisks*) and non-significant (*n.s.*) differences are highlighted. The control group (mCh-Cry2) is the same as in Fig. 6F as the experiments were performed in parallel. **D.** Representative western blot of lysates of HEK293 expressing either CDKL5-GFP (*left lane*) or GFP alone (*right lane*) to detect the levels of these two exogenous proteins (α-GFP immunoblotting, IB) as well as the endogenous CDKL5 target protein EB2, as such (α-EB2 IB) or phosphorylated at Ser222 (α-pEB2 IB). Note how EB2 is considerably more phosphorylated in CDKL5-GFP-expressing cells in comparison with what was observed in GFP-expressing cells, indicating that the exogenously expressed GFP-tagged kinase is catalytically active and enhances EB2 phosphorylation. **E.** As in *panel D*, for lysates of cells expressing GFP-tagged CDKL5 or one of the indicated disease-related mutants. Vinculin (vinc; α-vinculin IB), a housekeeping protein, was used as a lane loading control. **F.** Bar graph displaying the optical density analysis of western blotting experiments as shown in *panel E*. The columns report the ratio of the band intensities for pEB2 and EB2, normalized to the kinase (GFP) band intensity. Note how the relative phosphorylation level of EB2 is lower in cells expressing the two truncated kinases than in cells expressing the full-length CDKL5-GFP kinase.

These findings indicate that disease-related CTD truncations, by eliding the CTF and the most distal portion of the protein, can substantially alter the LLPS behavior of CDKL5 and, therefore, its subcellular localization.

Finally, we tested whether the observed impairment in LLPS caused by the disease-related truncations may also affect the catalytic function of CDKL5 in the cellular context. Towards this aim, we assessed the phosphorylation level of EB2, a known target of CDKL5 (Baltussen et al., 2018), in cells expressing either full-length CDKL5 or its two disease-related mutants (**Fig. 9D-F**).

As a preliminary step, we initially evaluated the ability of exogenously expressed CDKL5-GFP to enhance EB2 phosphorylation in HEK293 cells. Western blot analyses, using phospho-specific antibodies directed against EB2 phosphorylated at serine 222 (S222; ‘pEB2’), showed that CDKL5-GFP expression strongly enhanced EB2 phosphorylation relative to the GFP alone control condition (**Fig. 9D**).

Then, based on these findings, we tested whether the enhancement in EB2 phosphorylation induced by wild-type (*wt*) CDKL5-GFP was maintained or altered when expressing the disease-related mutants CDKL5-S726X or CDKL5-R781X. Towards this aim, we alternatively expressed these two GFP-tagged mutants in HEK293 cells to examine their relative ability to promote EB2 phosphorylation in comparison with full-length CDKL5-GFP (**Fig. 9E**).

This analysis showed that, when expressing either CDKL5 disease-related mutant, EB2 phosphorylation was significantly reduced as compared to what was observed when expressing *wt* CDKL5. A one-way ANOVA indicated overall significant differences between the experimental groups (F = 34.30, p < 0.001). Pairwise *post hoc* comparisons indicated significant differences between the CDKL5-GFP group and both the CDKL5-S726X and CDKL5-R781X groups (n = 4 in all groups, p < 0.001, Tukey’s *post hoc* test in both instances; **Fig. 9F**).

These results indicate that the very C-terminal portion of the protein, distal to a.a. 725, is crucial to ensure the physiological level of CDKL5 catalytic activity. This conclusion is also consistent with recent findings that missense mutations in the very distal part of the protein (T958R) can significantly alter its catalytic activity (Frasca et al., 2022).

Together with our previous observations, these findings show that the most C-terminal portion of CDKL5 (∼300 residues), encompassing the CTIF, has key functional roles in regulating both the subcellular localization of the protein, by means of LLPS, and its catalytic activity. As this region is elided by most disease-related truncating mutations, its loss –through the impairment of CDKL5 LLPS and functional activity– may play a key role in the molecular pathogenesis of CDD.

## DISCUSSION

The results of our experiments reveal the intrinsic ability of CDKL5 to form condensates through LLPS and identify its serine-rich CTD, and an internal fragment of it (CTIF), as key players in this process. Two disease-related mutations that elide the CTIF and the downstream tail of the CTD significantly impair both LLPS and the catalytic activity of CDKL5 in the cellular context. These findings uncover the role of LLPS in the functional biology of CDKL5 and in the molecular pathogenesis of CDD.

### LLPS of a nuclear, cytosolic, and synaptic kinase through a low-complexity serine-rich domain

Our morphological and functional experiments concurrently show that the intracellular foci formed by the CDKL5 protein display hallmark features of LLPS-driven condensates, including a rounded morphology, the ability to rapidly coalesce into larger droplets, a dynamic molecular exchange with the surrounding cellular environment, as revealed by FRAP, and the sensitivity to 1,6-Hex, which is known to disrupt the relatively weak molecular interactions underlying LLPS (Ulianov et al., 2021; Vaglietti et al., 2023). Moreover, optogenetic approaches and live-cell confocal imaging analyses using the optoDroplet system (Shin et al, 2017) further confirmed the ability of the protein CTD, and of the CTIF within it, to condensate in a rapid and reversible manner, another characteristic feature of LLPS-prone protein regions (Basu et al., 2020; Vaglietti et al., 2023).

Spontaneous and triggered condensation assays concurrently identified the CTD, and the CTIF, as key drivers of the phase separation process. The CTD alone did not display the condensation pattern of full-length CDKL5, thus indicating that the interplay between the CTD and the NTD defines the overall LLPS behavior of the entire protein. Indeed, the NTD was able to undergo a low degree of spontaneous condensation, consistent with recent observations that a catalytic domain of another protein kinase can form LLPS-driven condensates (Qiu et al., 2024).

Remarkably, the CTD and the CTIF contain a considerably high percentage (17.6% and 18%, respectively) of serine residues (8.01% on average in the human proteome). Serine-rich regions and polyserine (polyS) repeats have been implicated in protein oligo-/poly-merization, interactions, LLPS, and aggregation (Pelassa and Fiumara, 2015; Lilliu et al., 2018; Sabari et al., 2018; Marchetti et al 2021; Zhang et al., 2022), playing functional and evolutionary roles in biological entities ranging from viruses to multicellular organisms (Seifert et al. 2006; Yagi et al. 2014; Kim et al., 2016; Pelassa et al., 2019; Marchetti et al., 2021).

Our evolutionary analyses indicate that such a long serine-rich CTD in CDKL5 is phylogenetically ancient, as it is found even in species from older taxa along the vertebrate lineage (e.g. Agnatha), as well as in non-mammalian vertebrate clades. The serine richness of the CTD increased further in younger clades, reaching its highest level in metatherian and eutherian mammals, suggesting its functional relevance.

Based on sequence similarity, the vertebrate CDKL paralogs are split into two groups, one comprising CDKL1-4, the other CDKL5 alone (Canning et al., 2018). We found that the CDKL1–4 CTDs, which are considerably shorter than the CDKL5 CTD, are less enriched, or not enriched at all, in serine residues, displaying reduced or no LLPS propensity. This suggests the possibility that an ancestral vertebrate CDKL5 protein may have had a short CTD with some degree of serine enrichment and modest LLPS propensity. It is conceivable that, in the evolution of vertebrates, these two features were then greatly enhanced in CDKL5 and either maintained at a low level or entirely lost in its paralogs. These scenarios are consistent with the view that LCRs, including amino acid repeats (AARs), contribute to the functional diversification of paralog proteins and tend to be evolutionarily maintained if functionally relevant (Persi et al., 2016, 2023; Vaglietti et al., 2024).

CDKL genes have also been found in invertebrates. *C. elegans* and *D. melanogaster* possess only a single CDKL gene, more closely related to the vertebrate CDKL1-4 group, bearing a short CTD. Interestingly, the CTDs of both these two invertebrate CDKLs contain LCRs enriched in either A/Q (*D. melanogaster*) or N/S (*C. elegans*) residues, displaying some predicted LLPS propensity, in a configuration similar to that found in human CDKL4. These findings further support the notion that the presence of LLPS-prone LCRs may represent an ancient feature of CDKL genes that became particularly prominent in CDKL5. The compositionally distinct LCRs found in vertebrate and invertebrate CDKL CTDs (i.e. S-rich, A/Q-rich, and N/S-rich) may either derive from an ancestral LCR by divergent evolution or may have appeared independently in different clades by convergent evolution. Both these scenarios have been previously described for LCRs/AARs (Persi et al., 2016, 2023; Pelassa et al., 2019; Vaglietti et al., 2023, 2024).

Taken together, these evolutionary analyses strongly suggest that the peculiar compositional features of the CDKL5 CTD have been preserved and enhanced throughout phylogenesis because of a functional role, consistent with the growing evidence of the functionality of LCRs, including serine-rich/polyS sequences, in proteins.

Indeed, LCRs/AARs can mediate protein-protein interactions, oligo/-poly-merization, LLPS, and aggregation (Fiumara et al., 2010; Pelassa et al., 2014; Pelassa and Fiumara, 2015; Vaglietti and Fiumara, 2021; Vaglietti et al., 2023). Emerging evidence from our and other laboratories shows that polyS and serine-rich LCRs can drive protein LLPS and aggregation in physiological and pathological contexts (Lilliu et al., 2018; Sabari et al., 2018).

Consistent with these views, the results of our experiments directly implicate the serine-rich CTD, and its most distal portions, in the LLPS-driven condensation of CDKL5, as well as in the functional regulation of the protein catalytic activity. In principle, the serine-rich CTD may be functional in driving not only the condensation of CDKL5 alone, but also its co-condensation with other proteins into functional compartments (Ma et al., 2021; Vaglietti et al., 2023). Co-condensation with other proteins may underlie the recruitment of CDKL5 to subcellular compartments held together by LLPS-dependent mechanisms, such as the pre- and post-synaptic specializations, in which CDKL5 plays important physiological roles (Pizzo et al., 2016; Gurgone et al., 2023; Kontaxi et al., 2023). A few other protein kinases have been shown to undergo LLPS (López-Palacios and Andersen, 2023), indicating that condensation may represent a general physiological mechanism for the transient compartmentalization of kinase activity to specific subcellular domains.

Remarkably, serine-rich and polyS sequences are present in the LCRs of the prion-like CPEB2 and CPEB3 (Vaglietti et al., 2024) and other functional prions with roles in memory-related synaptic plasticity and protein-based inheritance in yeast (Si et al., 2003a,b; Theis et al., 2003; Crow et al., 2008; Fiumara et al., 2010, 2015; Tsvetkov et al., 2020). As these sequences can mediate LLPS-driven protein condensation (Sabari et al., 2018) as well as the formation of fibrillary structures (Lilliu et al., 2018), they may in principle promote the formation of more persistent forms of prion-like aggregates. Future studies may explore whether the serine-rich CTD may mediate, besides LLPS, also prion-like structural and functional switches of CDKL5 that may perpetuate its local activity within more persistent protein assemblies.

Overall, our results implicate the evolutionarily conserved serine-rich CTD, and its most distal portions, in the physiological LLPS-driven condensation of CDKL5, as well as in the functional regulation of the protein catalytic activity.

### LLPS in CDD and related neurodevelopmental disorders

The results of our study can help rationalize fundamental aspects of the molecular pathogenesis of CDD. Indeed, numerous disease-related mutations lead to the production of variably truncated forms of CDKL5 that lack, more or less completely, the CTD, which we have found to play a key role in LLPS. Our experiments, focusing on two of these mutations that elide one or two thirds of the CTIF, show that CDKL5 truncations can cause a significant reduction of the protein condensation capacity.

Recent observations show that a disease-related missense mutation (T958R) of one of the last few residues of CDKL5 significantly enhances its kinase function (Frasca et al., 2022), indicating how the most distal portion of the CTD can regulate the NTD catalytic activity. Remarkably, this region is absent in both the truncated mutants that we have studied, which display indeed similarly reduced LLPS and catalytic activity on the target protein EB2 in the cellular context.

Thus, by altering LLPS, truncations deleting the CTIF and the more distal portion of the protein significantly impair both CDKL5 localization and catalytic activity.

CDD has been initially considered a variant form of Rett syndrome (RTT), and both disorders, although distinct, display similarities at the phenotypic and clinical levels (Cutri-French et al., 2020). RTT is caused by mutations in the gene encoding the MeCP2 protein (Amir et al., 1999). Remarkably, MeCP2 has been shown to undergo LLPS and disease-related mutations significantly alter this process (Fan et al., 2020; Jiang et al., 2020; Wang et al., 2020). Thus, our findings that CDKL5 forms LLPS-driven condensates, and that this ability is impaired by disease-related truncating mutations, establish a striking parallelism at the molecular level between RTT and CDD.

LLPS and its disease-related dysregulation represent therefore key mechanisms underlying the function and dysfunction of two fundamental molecular players in closely related diseases. Ultimately, these findings indicate that LLPS dysregulation may represent a shared, fundamental mechanism in both CDKL5- and MeCP2-related neurodevelopmental disorders, opening new perspectives on their molecular pathogenesis.

## MATERIALS AND METHODS

### Bioinformatics

The primary sequences of the five paralog proteins of the human CDKL family (CDKL1-5) were derived from the Uniprot database (available at www.uniprot.org; IDs: Q00532, Q92772, Q8IVW4, Q5MAI5, O76039). These sequences were aligned using the Cobalt multiple sequence alignment tool (NCBI; available at https://www.ncbi.nlm.nih.gov/tools/cobalt/re_cobalt.cgi) and a graph displaying the degree of sequence conservation along the alignment was obtained (**Fig. 1D**). The primary sequences of CDKL5 ortholog proteins from 331 vertebrate species belonging to major vertebrate clades (Primates, Dermoptera, Scandentia, Glires, Laurasiatheria, Atlantogenata (i.e. Afrotheria + Xenarthra), Metatheria, Prototheria, Sauropsida, Amphibia, Actinopterygii, Chondrichthyes) and taxa (Agnatha; **Fig. 2A**) were obtained from NCBI (available at https://www.ncbi.nlm.nih.gov/protein/) and Uniprot databases. The list of IDs is reported in **Suppl. Table 1**. The Uniprot sequence IDs of the *Drosophila melanogaster* and *Caenorhabditis elegans* CDKL proteins are, respectively, Q9VMN3 and Q9U2H1. The phylogenetic tree of the species/clades of interest and the divergence times (million years ago, mya) between Primates and the other clades/taxa (i.e. Dermoptera, 79 mya; Scandentia, 85 mya; Glires, 87 mya; Laurasiatheria, 94 mya; Atlantogenata, 99 mya; Metatheria, 160 mya; Prototheria, 180 mya; Sauropsida, 319 mya; Amphibia, 352 mya; Osteichthyes, 429 mya; Chondrichthyes, 462 mya; Agnatha (*Petromyzon marinus*), 563 mya) were derived from TimeTree (Kumar et al., 2017; available at www.timetree.org; median divergence times were selected). The phylogenetic trees were downloaded in Newick format and further rearranged and processed using MegaX (Kumar et al., 2018).

Amino acid frequencies in human CDKL paralog and ortholog proteins, as well as their mean values across all proteins of the Uniprot reference human proteome, were calculated as in Marchetti et al. (2021) and Vaglietti et al. (2024). The CAST algorithm (Promponas et al., 2000) on the PlaToLoCo web server (Jarnot et al., 2020) was used to identify low-complexity regions in CDKL5, as shown in **Fig. 1A**. Thin bar graphs displaying the position of amino acids along protein primary sequences were generated using Excel. FuzDrop (Hardenberg et al., 2020) was used to calculate the overall LLPS propensity score (p_LLPS_) of proteins of interest and to obtain per-residue LLPS-propensity scores (P_DP_) along protein primary sequences. The disease-related CDKL5 variants that were analyzed (see **Fig. 8A**) were derived from the ClinVar database. These variants (related to the *Homo sapiens* cyclin dependent kinase like 5 (CDKL5), transcript variant III, mRNA; NCBI Reference Sequence: NM_001323289.2) included non-truncating (missense and in-frame indels) and truncating (nonsense and frameshift) mutations in the protein-coding part of the gene that were annotated as ‘pathogenic’ (**Suppl. Table 2**). An atomic level structural model of human CDKL5 was generated using the Colab AlphaFold2 software (Mirdita et al., 2022; available at https://colab.research.google.com/github/sokrypton/ColabFold/blob/main/AlphaFold2.ipynb), based on the primary sequence (Uniprot ID O76039). The model was downloaded as a file in PDB format. The available experimental structure of the kinase domain was derived from the PDB database (4bgq; Canning et al., 2018) in PDB format. These protein structures were visualized, processed, and pseudo-colored based on the FuzDrop per-residue P_DP_ score, or to identify specific residues, using the UCSF Chimera software (version 1.14; Meng et al., 2006).

### Plasmids and molecular cloning

Plasmid vectors for the expression of GFP- or mCh-Cry2-tagged proteins/peptides of interest were generated using the InFusion cloning system (Clontech-Takara). DNA fragments encoding CDKL5 or its fragments of interest were amplified by polymerase chain reacrion (PCR) from the pCAGIG vector containing the CDKL5 ORF (a kind gift of Dr. Alessandro Morellato, Molecular Biotechnology Center, University of Turin). The PCR products or interest were then subcloned into the pAcGFP-N1-In-Fusion-Ready plasmid (Clontech-Takara) or in the pX-mCh-Cry2 plasmid (Vaglietti et al., 2023).

In all cloning procedures, PCR products were obtained using the CloneAmp-HiFi PCR Premix (Clontech-Takara). The PCR primer oligonucleotides were designed with 15/18-bp extensions suitable for the In-fusion cloning system. A minimal Kozak sequence fragment (‘CACC’) was also included in the forward primer immediately upstream of the CDKL5 ATG start codon. An ATG start codon was also added after the CACC Kozak sequence for the expression of internal fragments of CDKL5. PCR products were separated by agarose gel electrophoresis and purified from agarose gels using the NucleoSpin kit (Macherey-Nagel). Purified PCR products were subcloned into linearized plasmid vectors of interest using the In-fusion cloning system following the manufacturer’s protocol. Plasmids were amplified using Stellar chemically competent cells (Clontech-Takara) and purified using the NucleoSpin plasmid Mini kit (Macherey-Nagel) or the NucleoBond Xtra MIDI EF kit (Macherey-Nagel). For all the plasmids that were generated, the protein coding sequences were verified by Sanger sequencing.

Using this general approach, we generated plasmids for the expression of GFP-tagged CDKL5, its fragments of interest, i.e. the N-terminal domain (NTD, a.a. 1-297) and the C-terminal domain (CTD, a.a. 278-960), two C-terminally truncated disease-related mutants, i.e. CDKL5-Δ726-960 (a.a. 1-725) and CDKL5-Δ781-960 (a.a. 1-780), and an internal deletion mutant (CDKL5-ΔCTIF), lacking a.a. 657-840. The latter mutant was obtained through a multi-fragment In-Fusion cloning reaction with two PCR products of the ORF fragments, encoding a.a. 1-656 and a.a. 841-960, that were fused together at one of their ends and then to the vector.

To generate suitable plasmid vectors for optoDroplet induction experiments (Shin et al., 2017), the nucleotide sequences encoding CDKL5 fragments of interest, i.e. the NTD, the CTD, the C-terminal fragments (a.a. 726-960 and a.a. 781-960), and the internal CTIF fragment (a.a. 657-840) were PCR-amplified from the pAcGFP-N1-CDKL5 plasmid using the CloneAmp-HiFi PCR Premix with primers bearing suitable 15-bp extensions to allow their subcloning into the pX-mCh-Cry2 plasmid (Vaglietti et al., 2023) using the In-Fusion cloning system. These vectors allowed the expression of the CDKL5 fragments of interest as fusion proteins with mCherry (mCh), for fluorescence visualization, and a fragment of the cryptochrome 2 protein of *A. thaliana* (Cry2), which can trigger the condensation of LLPS-prone peptides in a blue light-dependent manner (Shin et al., 2017).

### Cell culture and neuronal differentiation

NG108-15 cells (Nelson et al., 1976; Christian et al., 1977; Han et al., 1991) were maintained in Dulbecco’s Modified Eagle’s Medium (DMEM; Sigma) with 10% fetal bovine serum (FBS; Sigma-Aldrich) and 1X HAT supplement (ThermoFisher), 2mM L-glutamine (Sigma-Aldrich), 100 units/mL penicillin (Sigma-Aldrich), 100 µg/mL streptomycin (Sigma-Aldrich), as in Mitchell et al. (2007). To induce neuronal differentiation, NG1081-15 cells were plated onto poly-D-lysine-coated glass coverslips (0.1 mg/mL, Sigma) in a 24-well plate and, after 24-48 h, the culture medium was replaced with a differentiation medium consisting of DMEM with 1% FBS, 1X HAT, 1 mM dibutyryl-cAMP (dbcAMP; Sigma), 2mM L-glutamine (Sigma-Aldrich), 100 units/mL penicillin (Sigma-Aldrich), 100 µg/mL streptomycin (Sigma-Aldrich), for 7-14 days (Mitchell et al., 2007). Half of the differentiation medium in the culture wells was replaced with fresh one on alternate days. HEK293 cells (293-F strain; Thermo Fisher) were maintained following standard procedures in DMEM (Thermo Fisher) supplemented with 10% fetal bovine serum (FBS), 2mM L-glutamine (Sigma-Aldrich), 100 units/mL penicillin (Sigma-Aldrich), 100 µg/mL streptomycin (Sigma-Aldrich) in an atmosphere with 5% CO2 at 37 °C. The cells that were used for immunocytochemistry or plasmid transfection were plated onto glass coverslips coated with poly-L-lysine (0.1 mg/mL, Sigma-Aldrich) in 24-well plates. For optoDroplet experiments, HEK293 cells were plated into 50 mm plastic culture dishes with a central 14 mm glass coverslip (Mattek) coated with poly-L-lysine.

Hippocampal cell cultures were prepared from P18 embryos (400-800 cells/mm^2^) as in Russo et al. (2017). Immunocitochemistry experiments on hippocampal neurons were performed on DIV13– 15.

### Cell culture transfection

Transfections of NG108-15 and HEK293 cells were performed 24 h after plating using Lipofectamine 2000 (ThermoFisher) following the manufacturer’s protocol, using 0.25-1 μg of plasmid DNA for each coverslip in a single well of a 24-well plate with a 3:1 Lipofectamine (μl) to DNA (μg) ratio. When transfecting NG108-15 cells, the culture medium containing the transfection mix was replaced four hours after transfection with fresh medium. Twenty-four hours after transfection, the culture medium was replaced with the differentiation culture medium and NG108-15 cells were allowed to differentiate for 7-14 days before fixation. For optoDroplet experiments, the transfection of HEK293 cells was performed on non-adherent cells in suspension, immediately prior to plating, using the same reagents (Vaglietti et al., 2023). The cells were then plated onto poly-L-lysine coated coverslips. HEK293 cells were fixed at 24, 48 or 72 h after transfection. Both NG108-15 and HEK293 cells were fixed, after rinsing with phosphate-buffered saline (PBS; 3x), using 4% paraformaldehyde (PFA) in PBS (pH 7.4) for 15-20 min at RT. After rinsing in PBS (3x), the coverslips were mounted onto microscope slides using DAKO mounting medium (Agilent).

### Immunocytochemistry (ICC)

Hippocampal, NG108-15, and HEK293 cells growing on glass coverslips were fixed in 4% PFA after rinsing with PBS (3x). The cells were then permeabilized using 0.1% Triton X-100 in PBS, pH 7.4, for 15 minutes, rinsed in PBS (3x), and then incubated for 1 h in blocking solution (1% bovine serum albumin, BSA; ThermoFisher) in PBS). The cells were then incubated with primary antibodies against human CDKL5 (Santa Cruz Biotechnology, sc-376314, dilution 1:50 in 0.1% BSA in PBS) for 3 hours at room temperature (RT). After rinsing with PBS (3x), the cells were incubated with an Alexa Fluor 488-conjugated secondary antibody (Thermo Fisher Cab#A-11001, dilution 1-1000) for 1 hour at RT and rinsed with PBS (3x). Finally, nuclei were stained with 300 nM DAPI (ThermoFisher) in PBS for 5 minutes at RT. In some experiments, filamentous actin was stained using the Phalloidin CruzFluor 555 conjugate (Santa Cruz Biotechnology, sc-363794) for 90 minutes at RT and rinsed in PBS (3x). The coverslips were mounted onto microscope slides using DAKO mounting medium (Agilent).

### Confocal and super-resolution (Airyscan) fluorescence microscopy

Confocal and super-resolution imaging (Airyscan) of hippocampal, NG108-15, and HEK293 cells expressing fluorescently-labelled proteins of interest, or stained otherwise with secondary fluorescent antibodies, fluorescent phalloidin, and/or DAPI, was performed using a ZEISS LSM 800 Airyscan microscope (Zeiss) with a 63X immersion objective using default Airyscan settings. Some confocal images were acquired using a Fluoview FV300 confocal microscope (Olympus). Fluorescence imaging was also performed in some experiments using the Mica Microhub imaging platform (Leica Microsystems) and Thunder deconvolution using the LasX software platform (Leica Microsystems). Image analysis was performed using FIJI-ImageJ (Schindelin et al., 2012).

### Liquid-liquid phase separation (LLPS) analysis

The spontaneous formation of condensates by GFP-tagged CDKL5, its fragments, and its disease-related mutants, was quantified in confocal fluorescence images of HEK293 and NG108-15 cells acquired at 24-72 h after transfection. *Z*-stack maximum intensity projections of fluorescence confocal images of 233×233 µm, or 350×350 µm, microscopy fields were converted into 8-bit images using FIJI/ImageJ (NIH). For each image, brightness and contrast were adjusted alternately to highlight either the fluorescent condensates alone or the entire profiles of fluorescent nuclei/cells. These two adjusted images were used to automatically quantify, for each image, the area and number of individual condensates, the total area occupied by condensates, and the total area of fluorescent cells/nuclei using *ad hoc* CellProfiler (Carpenter et al., 2006) pipelines. From these parameters, we calculated for each image the proportion of the total cell/nuclei fluorescent area occupied by condensates and the mean size of each condensate. For each experimental group, we analyzed 10-40 microscopy fields from cell cultures derived from at least three independent biological replicates of the same experiment.

To quantify rapid LLPS induction, and its kinetics, using the optoDroplet system (Shin et al.,2017), HEK293 cells, grown onto poly-L-lysine-coated glass coverslips and expressing for 24-48 h either CDKL5 fragments of interest fused to mCh-Cry2, or mCh-Cry2 alone, were photoactivated using 488 nm blue light emitted by an LED light array within a custom-made apparatus, with a working distance of 10 cm between the LEDs and the cell culture, for 4–5 minutes. (Vaglietti et al., 2023). In pilot experiments, the blue light intensity was set at a level that triggered the condensation of an LLPS-prone mutant form of mCh-Cry2 (mCh-Cry2Olig, used as a positive control) but not of mCh-Cry2 (Shin et al., 2017). Two minutes after photoactivation, the cells were rinsed in PBS and fixed with 4% PAF in PBS. After rinsing in PBS (3x), coverslips were mounted onto microscope slides for subsequent fluorescent confocal imaging to detect the mCherry red fluorescence. Control cells were treated in the same manner, except that photoactivation was omitted. LLPS-driven condensate formation was quantified, as described above, by calculating, for each construct and treatment, the relative proportion of the mCh-positive cell area occupied by condensates using an *ad hoc* CellProfiler pipeline.

### Live cell imaging

Live-cell imaging experiments of HEK293 cells expressing GFP- or mCh-tagged constructs of interest were performed using a Fluoview FV300 confocal microscope (Olympus). For these experiments the culture medium was replaced with Hank’s balanced saline solution (HBSS) and cells were incubated in a custom-made on-stage incubator for temperature control (Ragazzini et al., 2018). To highlight the coalescence of CDKL5-GFP condensated, cell cultures were imaged every 10 s. To study the condensation kinetics of the CDKL5 CTIF fragment using the optoDroplet system (see above), HEK293 cells expressing CTIF-mCh-Cry2 were imaged 30 s before and 10 min after (one image every 30 s) a photoactivating pulse delivered using the 488 nm laser light (as in Vaglietti et al., 2023). Images were further analyzed using FIJI software and CellProfiler, as described above.

### 1,6-Hexanediol (1,6-Hex) treatments

To define the sensitivity to 1,6-Hex of intracellular condensates formed by endogenous CDKL5, HEK293 cultures were exposed to a final concentration of 5% 1,6-Hex in culture medium for 5 or 15 minutes (Ulianov et al., 2021; Vaglietti et al., 2023). The cells were then rinsed with PBS (3x) and fixed in 4% PFA in PBS (15-20 min at RT) and processed for ICC using a primary anti-CDKL5 antibody (Santa Cruz Biotechnology, sc-376314; see above). The treatment with 1,6-Hex was omitted in control cultures. The proportion of the fluorescent cell area occupied by condensates was measured as described above. The same protocol was also used to quantify the sensitivity to 1,6-Hex of condensates formed by exogenously expressed CDKL5-GFP at 24-48 hours after transfection.

### Fluorescence recovery after photobleaching (FRAP)

To perform FRAP experiments, HEK293 cells were plated at low density in 8-well glass bottom microslides (IBIDI) and transfected to express CDKL5-GFP. FRAP experiments were performed 24–48 h after transfection, on a TCS SP8 confocal microscope (Leica) equipped with an HCX PL APO 63X/1.4 oil objective, an adaptive focus control, and stage incubator (OkoLab) to maintain temperature at 37 °C in a 5% CO_2_ atmosphere. Each individual cell nucleus was imaged for 5 iterations (5X zoom, pinhole 5AU) and then then a CDKL5-GFP condensate contained within a 3 µm^2^ circular region of interest (ROI) was bleached using high (100%) laser power at 488 nm (Argon laser at 70% initial power, 1 iteration). After bleaching, images were taken every 1.625 sec to monitor fluorescence recovery. For each condensate that was studied, fluorescence intensities measured before and after photobleaching were quantified for the photobleached region of interest or ‘ROI’ (ROI1), a second ROI encompassing the entire cell (ROI2), and a third ROI encompassing the non-fluorescent background region (ROI3) using the LasX software (Leica). The data were analyzed using the EasyFrap tool (Kolouras et al., 2018) by subtracting background fluorescence (as detected in ROI3) and compensating for imaging-related fluorescence decay (as detected in ROI2). The final FRAP curve was obtained from EasyFRAP using double normalization and was fitted to a double exponential curve to calculate the mobile fraction (*Mf*) and the recovery half-time (*t_1/2_*).

### Correlative light and electron microscopy (CLEM)

HEK293 cells, plated onto 35-mm dishes containing a gridded glass coverslip (MatTek) and expressing CDKL5-GFP for 24-48 h, were fixed with 4% PFA, 1% glutaraldehyde in 0.1 M HEPES buffer (pH 7.4). To obtain additional reference points for the precise alignment of the fluorescence and EM images, cell cultures expressing CDKL5-GFP were also stained with MitoTracker Red CMXRos (200 nM, ThermoFisher) and Hoechst 33342 (ThermoFisher) to label mitochondria and cell nuclei, respectively. Confocal images of cells expressing CDKL5-GFP were acquired using a FluoVIEW 3000 RS system (Evident Scientific) equipped with a UPLSAPO 60X/1.3 silicon objective. Z-stacks encompassing the entire cell were acquired with an optical section depth of 0.21 μm and the images were deconvolved using Huygens Software (SVI) prior to alignment with corresponding EM images. The cell cultures were postfixed with reduced osmium (1% OsO4, 1.5% potassium ferrocyanide in 0.1 M cacodylate buffer, pH 7.4) for 1 h on ice. After multiple washes in ultrapure water, they were incubated in 0.5% uranyl acetate overnight at 4°C. Samples were then dehydrated with increasing ethanol concentrations, embedded in epoxy resin, and polymerized for 48 h at 60 °C. Ultrathin serial sections (70 nm thick) were obtained using a UC7 ultramicrotome (Leica), collected on formvar-carbon coated copper slot grids, stained with uranyl acetate and Sato’s lead solutions, and observed under a Talos L120C transmission electron microscope (ThermoFisher) operating at 120 kV. Images were acquired using a Ceta CCD camera (ThermoFisher) and aligned with confocal images using the Fiji macro-plugin Big Warp.

### Cell lysate preparation

HEK293 cell cultures expressing either CDKL5-GFP, CDKL5-S726X-GFP, CDKL5-R781X-GFP, or GFP alone, for 24 h were lysed using an ice-cold lysis buffer (150 mm NaCl, 50 mm Tris, 5% glycerol, 1% NP-40, 1 mM MgCl2) supplemented with 200 mM sodium orthovanadate, 100 mM dithiothreitol (DTT), 100 mM phenylmethylsulfonyl fluoride (PMSF), 1M sodium fluoride and protease inhibitors (SIGMAFAST protease inhibitor cocktail tablets, EDTA-free). The lysates were centrifuged at 13,000 g for 10 minutes at 4 °C to remove cell debris, and the supernatants were stored at -80 °C until used for Western blot experiments. Sample protein concentrations were determined using a Nanodrop spectrophotometer (ThermoFisher).

### Western blotting

Western blotting experiments have been performed as in Gurgone et al. (2023). Briefly, cell lysates were boiled in SDS sample buffer, separated by SDS–PAGE, and the proteins were then blotted to a PVDF membrane following standard procedures. Next, PVDF membranes were blocked in 5% non-fat dry milk in TBS with 0.1% Tween 20 for 1 h at RT and incubated with primary antibodies overnight at 4°C. We used primary antibodies directed against vinculin (ImmunologicalSciences, cat. # AB-81457, 1:1000 dilution), EB2 (Abcam cat. # ab45767, 1:1000 dilution), phosphorylated EB2 (at serine 222; Covalab, cat. # 00117741, 1:1000 dilution) and GFP (ChromoTek, cat. # pabg1-100, 1:1000 dilution). After rinsing in 0.1% Tween 20 in TBS, the membranes were incubated with the appropriate secondary antibodies conjugated with horseradish peroxidase (HRP; anti-rat or anti-rabbit, 1:5000 dilution; Sigma-Aldrich) for 1 h at RT. The chemiluminescence signal was visualized using the Clarity Western ECL Blotting Substrate (Bio-Rad; Italy), acquired with a ChemiDoc imager (Bio-Rad), and analysed using Image-J software (NIH). Band intensity values related to anti-GFP and anti-EB2 antibodies were initially normalized, in each lane, to the intensity of the band obtained using the anti-vinculin antibody. As detailed in the *Results* section, the intensity of the band obtained with the phosphospecific anti-EB2 antibody was normalized to that of the band obtained with the non-phosphospecific anti-EB2 antibody. The normalized intensity was further normalized to the intensity of the band obtained with the anti-GFP antibody, which reflects the expression level of the exogenously expressed kinase (wild-type or mutant) in each lysate.

### Statistics

Data and statistical analyses were performed using Excel (Microsoft) and Statistica (TIBCO) software. Data are expressed as mean ± standard error of mean (SEM). Student t-test, ANOVA (one- or two-way) followed by *post hoc* tests, and Fisher’s exact test were performed, when appropriate, to assess statistical significance. A p-value < 0.05 was considered statistically significant.

## ACKNOWLEDGEMENTS

This research was supported by RiLo2022/2023 grants from the University of Turin to F.F.

## SUPPLEMENTARY MATERIALS

*-Suppl. Figures 1-4* with legends.

*-Suppl. Tables 1-2*.

