## Supplementary Materials for "The CDKL5 kinase undergoes liquid-liquid phase separation driven by a serine-rich C-terminal region and impaired by neurodevelopmental disease-related truncations"

### SUPPLEMENTARY FIGURE LEGENDS

**Suppl. Figure 1** – *Differential distribution of amino acids in the NTD and CTD of CDKL5*

**A**. Schematic representation of the CDKL5 protein with the distribution of the 20 amino acids (indicated by single letter codes, form A to Y), represented by *thin vertical red lines* along its primary sequence. The NTD is highlighted in *cyan*, the CTD in *turquoise*. Note how, certain residues are more concentrated in one of the two domains, like serine (S) in the CTD and leucine (L) in the NTD, while other residues, like alanine (A), are more evenly distributed. **B.** The graphs in the *left column* report the percent of the 20 residues in the CDKL5 full-length (FL) protein (*upper panel*), in the NTD (*middle panel*) and in the CTD (*lower panel*). The bar graphs in the *middle column* plot the difference between the percent occurrence of each amino acids in the entire CDKL5 sequence (FL), or in the NTD and CTD domains, and its mean percent occurrence across all proteins of the human proteome. The graphs in the *right column* are as those of the *middle column* but for classes of amino acids. *Hydrophobic residues*: small (*sma*; A, G), aliphatic (*ali*; V, L, I), aromatic (*aro*; F, W, Y), and sulphurated (*sul*; M, C). *Polar* residues (*polar*): N, Q, S. *Negatively charged* (-): D, E. *Positively charged* residues (+): H, K, R. P is the only cyclic (*cy*.) residue. **C.** AlphaFold structural model of the entire human CDKL5 protein (*upper panel*) showing the high concentration of leucine residues (L, highlighted in *cyan*) in the α-helical structures of the NTD. The *lower panel* highlights the positions of leucine residues on the available experimental structure of the NTD (PDB: 4bgq; Canning et al., 2018).

**Suppl. Figure 2** – *CDKL orthologs of* Caenorhabditis elegans *and* Drosophila melanogaster *have C-terminal LCRs enriched in Q/A or N/S residueszz and predicted to undergo LLPS.*

**A**. In the *upper panel*, plot of the per-residue FuzDrop LLPS propensity score ($P_{DP}$ score) along the *Caenorhabditis elegans* CDK-like kinase primary sequence. The *black* peaks highlight the protein regions with the highest LLPS propensity ($P_{DP}$ score > 0.6). The *white bar* above represents the protein with asparagine (N) and serine (S) residues highlighted in *red.* Note how the protein regions predicted to undergo LLPS correspond to the C-terminal N/S-rich tail. The overall protein pLLPS score (0.22) predicted by FuzDrop is reported *on the left*. The *middle panel* shows the per-residue $P_{DP}$ score along the *Drosophila melanogaster* CDK-like kinase primary sequence and the *white bar* above represents the protein with alanine (A) and glutamine (Q) residues highlighted in *red*. Even in this case, the largest predicted LLPS-prone region comprises the A/Q-rich portion of the protein. *Below*, the same graph shown in *Fig 3A* for human CDKL5 is reported here for comparison.

**Suppl. Figure 3** – *CDKL5 displays the longest serine-rich CTD among the members of the human CDKL protein family*

**A.** Per-residues FuzDrop LLPS propensity score ($P_{DP}$ score) along the primary sequences of the five human CDKL proteins (CDKL1-5). On the *left* of each graph, the overall FuzDrop pLLPS score of the protein is reported. Note how only CDKL5 displays a strong overall propensity to undergo LLPS (pLLPS = 0.99). Of the other paralog proteins, only CDKL3, which has an intermediate length CTD, displayed a borderline pLLPS score of 0.59. However, CDKL3, and to a lesser extent CDKL2 and CDKL4, display some peaks in the prediction profile that are clearly above-threshold (highlighted in *black*), which may suggest that some regions of these proteins have some degree of LLPS propensity. **B.** The *left* bar graph displays the FuzDrop LLPS propensity score (pLLPS score) of human CDKL1-5 primary sequences. The *middle* bar graph displays the percent occurrence of serine residues in human CDKL1-5 primary sequences. The scatterplot on the *right* displasy the significant correlation between the pLLPS score and the percent occurrence of serine residues for the five members of the CDKL protein family ($r = 0.95$, $n = 5$, $p<0.02$).

**Suppl. Figure 4** – *Truncating and non-truncating pathogenic mutations differentially affect the NTD and CTD of CDKL5*

**A.** Bar graph showing the proportion (%) of non-truncating (NTMs, either missense or inframe indel) and truncating (TMs, either nonsense or frameshift) pathogenic mutations (as reported in the *ClinVar* database) affecting the CDKL5 NTD (*cyan*) or CTD (*turquoise*). The distribution of the two mutation classes across the two domains is significantly different ($p<0.01$, Fisher's exact test). NTMs affect almost exclusively the NTD, whereas TMs affect both domains.

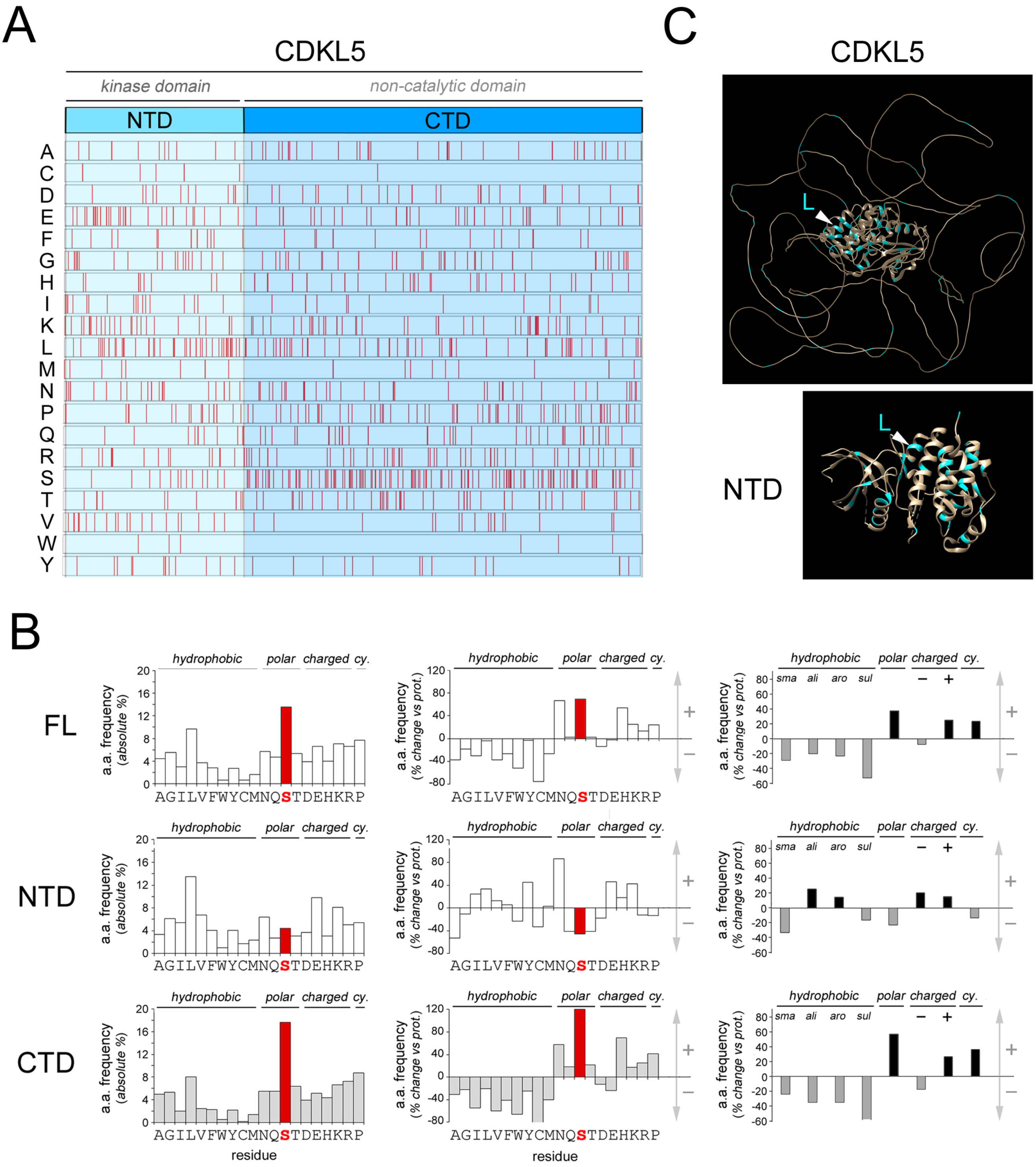


Suppl. Figure 1

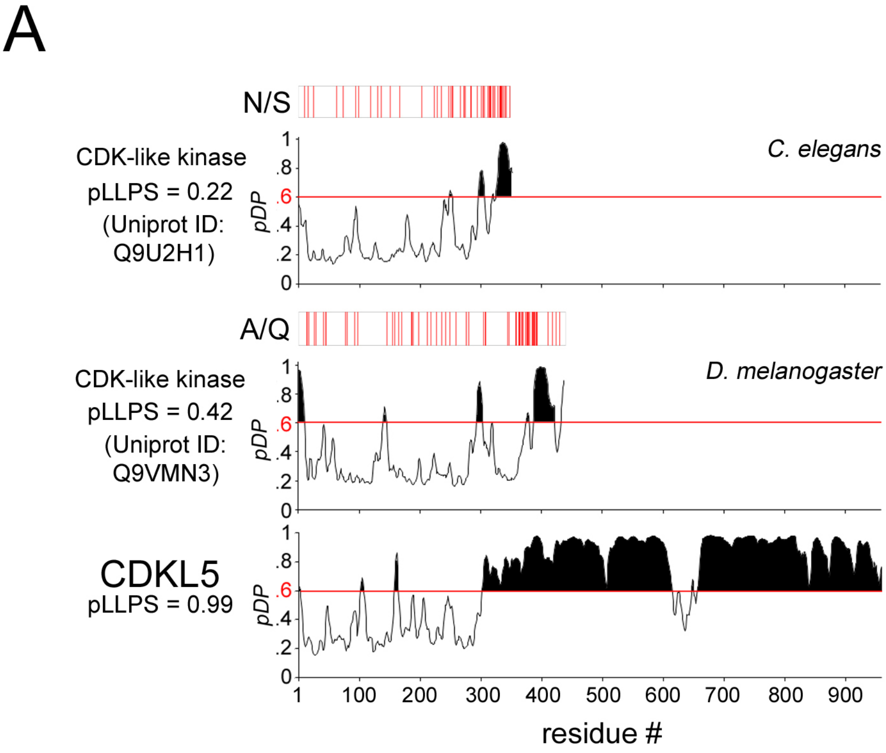


Suppl. Figure 2

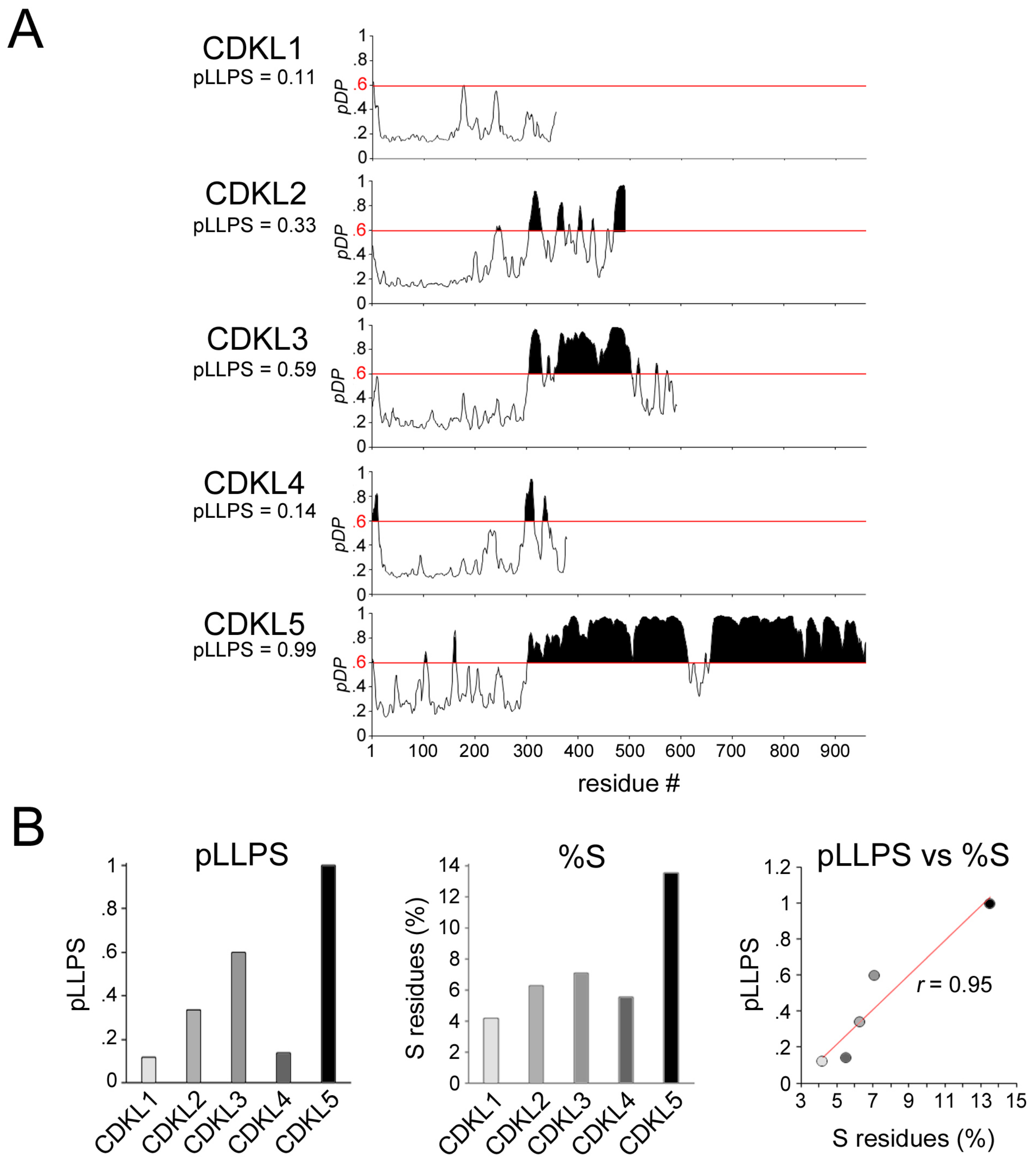


Suppl. Figure 3

A

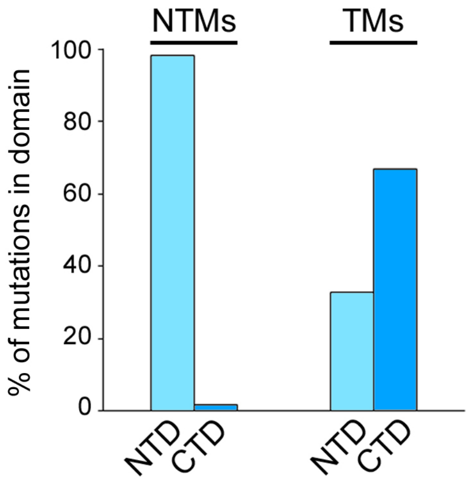


Suppl. Figure 4

| **Suppl. Table 1** - IDs of CDKL5 ortholog protein sequences | |
|---|---|
| **Species** | **Sequence ID (NCBI/Uniprot)** |
| *Homo sapiens* | NP_001310218.1 |
| *Pan paniscus* | XP_014198883.1 |
| *Pongo abelii* | XP_024096718.1 |
| *Nomascus leucogenys* | XP_003261162.1 |
| *Hylobates moloch* | XP_032612543.1 |
| *Macaca mulatta* | XP_028698065.1 |
| *Macaca fascicularis* | A0A7N9IGG3 |
| *Cercocebus atys* | XP_011891136.1 |
| *Chlorocebus sabaeus* | XP_007989371.2 |
| *Aotus nancymaae* | XP_012289642.1 |
| *Callithrix jacchus* | XP_035144933.1 |
| *Sapajus apella* | XP_032141915.1 |
| *Saimiri boliviensis boliviensis* | XP_010333196.1 |
| *Carlito syrichta* | XP_008063660.1 |
| *Microcebus murinus* | XP_012640148.1 |
| *Propithecus coquereli* | XP_012507129.1 |
| *Prolemur simus* | A0A8C8ZDJ8 |
| *Otolemur garnettii* | XP_003791925.1 |
| *Galeopterus variegatus* | XP_008578054.1 |
| *Tupaia chinensis* | XP_027631275.1 |
| *Mus musculus* | A0A0G2JGW6 |
| *Mus caroli* | XP_021009263.1 |
| *Mus pahari* | XP_029390005.1 |
| *Mastomys coucha* | XP_031226261.1 |
| *Arvicanthis niloticus* | XP_034341733.1 |
| *Rattus norvegicus* | XP_017457780.1 |
| *Microtus oregoni* | XP_041498332.1 |
| *Microtus ochrogaster* | XP_013206022.1 |
| *Arvicola amphibius* | XP_038172589.1 |
| *Mesocricetus auratus* | XP_005074284.1 |
| *Cricetulus griseus* | XP_027290329.1 |
| *Peromyscus maniculatus bairdii* | XP_015847608.1 |
| *Peromyscus leucopus* | XP_028729097.1 |
| *Onychomys torridus* | XP_036029834.1 |
| *Nannospalax galili* | XP_008833908.1 |
| *Jaculus jaculus* | XP_004667977.1 |
| *Dipodomys ordii* | XP_012887583.1 |
| *Octodon degus* | XP_023575351.1 |
| *Chinchilla lanigera* | XP_005414499.1 |
| *Cavia porcellus* | XP_023419382.1 |
| *Heterocephalus glaber* | XP_004866202.1 |
| *Fukomys damarensis* | XP_010641954.1 |
| *Ictidomys tridecemlineatus* | XP_005341193.1 |
| *Urocitellus parryii* | XP_026252522.1 |
| *Marmota marmota marmota* | XP_015343550.1 |

| | |
|---|---|
| *Sciurus vulgaris* | A0A8D2E472 |
| *Ochotona princeps* | XP_004597541.1 |
| *Ochotona curzoniae* | XP_040853602.1 |
| *Oryctolagus cuniculus* | XP_008270622.1 |
| *Eumetopias jubatus* | XP_027978412.1 |
| *Zalophus californianus* | XP_027463768.1 |
| *Callorhinus ursinus* | XP_025713447.1 |
| *Odobenus rosmarus divergens* | XP_004401557.1 |
| *Neomonachus schauinslandi* | XP_021539522.1 |
| *Mirounga leonina* | XP_034856219.1 |
| *Phoca vitulina* | XP_032245536.1 |
| *Enhydra lutris kenyoni* | XP_022363620.1 |
| *Lontra canadensis* | XP_032696727.1 |
| *Mustela putorius furo* | XP_004758990.1 |
| *Mustela erminea* | XP_032185542.1 |
| *Ursus maritimus* | XP_040489717.1 |
| *Ursus arctos horribilis* | XP_026336015.1 |
| *Ailuropoda melanoleuca* | XP_034505588.1 |
| *Vulpes lagopus* | XP_041597526.1 |
| *Vulpes vulpes* | XP_025853281.1 |
| *Canis lupus familiaris* | XP_038378906.1 |
| *Lynx pardinus* | A0A485PK05 |
| *Lynx canadensis* | XP_030161412.1 |
| *Puma concolor* | XP_025790422.1 |
| *Acinonyx jubatus* | XP_026910444.1 |
| *Felis catus* | XP_011289926.1 |
| *Panthera pardus* | XP_019287177.1 |
| *Panthera leo* | A0A8C8XQ28 |
| *Hyaena hyaena* | XP_039103298.1 |
| *Suricata suricatta* | XP_029786004.1 |
| *Manis pentadactyla* | XP_036768773.1 |
| *Equus caballus* | F7BV09 |
| *Equus asinus asinus* | A0A8C4MX80 |
| *Tursiops truncatus* | XP_033705291.1 |
| *Globicephala melas* | XP_030699571.1 |
| *Lagenorhynchus obliquidens* | XP_026955487.1 |
| *Orcinus orca* | XP_004269686.1 |
| *Delphinapterus leucas* | XP_022421190.1 |
| *Monodon monoceros* | XP_029096414.1 |
| *Phocoena sinus* | XP_032475235.1 |
| *Lipotes vexillifer* | XP_007467463.1 |
| *Physeter catodon* | XP_023971034.1 |
| *Balaenoptera acutorostrata scammoni* | XP_007173621.1 |
| *Balaenoptera musculus* | XP_036696585.1 |
| *Bos taurus* | XP_024844263.1 |
| *Bos indicus* | XP_019811075.1 |
| *Bos mutus* | XP_005890072.1 |
| *Bubalus bubalis* | XP_006042021.1 |

| | |
|---|---|
| *Ovis aries* | XP_027818684.1 |
| *Capra hircus* | XP_017899375.1 |
| *Oryx dammah* | XP_040122478.1 |
| *Odocoileus virginianus texanus* | XP_020746919.1 |
| *Catagonus wagneri* | A0A8C3WMX6 |
| *Sus scrofa* | XP_020936211.1 |
| *Camelus bactrianus* | XP_010952871.1 |
| *Vicugna pacos* | XP_006217756.1 |
| *Myotis myotis* | XP_036160929.1 |
| *Myotis davidii* | XP_006765193.1 |
| *Myotis lucifugus* | XP_006088613.1 |
| *Pipistrellus kuhlii* | XP_036295483.1 |
| *Eptesicus fuscus* | XP_008152773.1 |
| *Miniopterus natalensis* | XP_016054832.1 |
| *Molossus molossus* | XP_036126855.1 |
| *Artibeus jamaicensis* | XP_037013391.1 |
| *Phyllostomus discolor* | XP_028378889.1 |
| *Desmodus rotundus* | XP_024425648.1 |
| *Pteropus giganteus* | XP_039708257.1 |
| *Pteropus vampyrus* | XP_011361303.1 |
| *Pteropus alecto* | XP_006911691.1 |
| *Rousettus aegyptiacus* | XP_036082329.1 |
| *Hipposideros armiger* | XP_019521305.1 |
| *Rhinolophus ferrumequinum* | XP_032975742.1 |
| *Sorex araneus* | XP_004612276.1 |
| *Erinaceus europaeus* | XP_007538805.1 |
| *Echinops telfairi* | XP_004709992.1 |
| *Chrysochloris asiatica* | XP_006835657.1 |
| *Orycteropus afer afer* | XP_007956575.1 |
| *Loxodonta africana* | XP_003416076.1 |
| *Trichechus manatus latirostris* | XP_004382879.1 |
| *Choloepus didactylus* | XP_037678788.1 |
| *Dasypus novemcinctus* | XP_023442279.1 |
| *Vombatus ursinus* | XP_027691207.1 |
| *Phascolarctos cinereus* | XP_020863051.1 |
| *Trichosurus vulpecula* | XP_036599306.1 |
| *Sarcophilus harrisii* | XP_031814781.1 |
| *Monodelphis domestica* | XP_001380717.1 |
| *Tachyglossus aculeatus* | XP_038612960.1 |
| *Ornithorhynchus anatinus* | XP_007668543.1 |
| *Zonotrichia albicollis* | A0A8D2NI83 |
| *Junco hyemalis* | A0A8C5J041 |
| *Molothrus ater* | XP_036243363.1 |
| *Serinus canaria* | XP_009087940.1 |
| *Motacilla alba alba* | XP_038014644.1 |
| *Passer montanus* | XP_039570224.1 |
| *Pyrgilauda ruficollis* | XP_041328436.1 |
| *Corvus moneduloides* | XP_031956642.1 |

| | |
|---|---|
| *Corvus kubaryi* | XP_041892981.1 |
| *Corvus cornix cornix* | XP_039407037.1 |
| *Corvus brachyrhynchos* | XP_017583950.1 |
| *Taeniopygia guttata* | XP_030137515.3 |
| *Lonchura striata domestica* | XP_031359558.1 |
| *Catharus ustulatus* | XP_032908699.1 |
| *Ficedula albicollis* | XP_016154268.1 |
| *Sturnus vulgaris* | XP_014731290.1 |
| *Cyanistes caeruleus* | XP_023774212.1 |
| *Parus major* | XP_015486002.1 |
| *Pseudopodoces humilis* | XP_014119003.1 |
| *Cyanoderma ruficeps* | A0A8C3QSC8 |
| *Zosterops lateralis melanops* | A0A8D2PQL6 |
| *Hirundo rustica* | XP_039947200.1 |
| *Pipra filicauda* | XP_027594058.2 |
| *Manacus vitellinus* | XP_008920602.2 |
| *Lepidothrix coronata* | XP_017667778.1 |
| *Neopelma chrysocephalum* | XP_027544700.1 |
| *Empidonax traillii* | XP_027759399.1 |
| *Acanthisitta chloris* | XP_009080855.1 |
| *Corapipo altera* | XP_027493816.1 |
| *Chiroxiphia lanceolata* | XP_032534463.1 |
| *Nestor notabilis* | XP_010013824.1 |
| *Strigops habroptila* | XP_030332563.1 |
| *Melopsittacus undulatus* | A0A8C6IZ63 |
| *Falco cherrug* | XP_027666649.1 |
| *Falco rusticolus* | XP_037233151.1 |
| *Falco peregrinus* | XP_027634764.1 |
| *Falco tinnunculus* | A0A8C4V4U7 |
| *Falco naumanni* | XP_040438571.1 |
| *Otus sunia* | A0A8C8AJA3 |
| *Bubo bubo* | A0A8C0F0W5 |
| *Athene cunicularia* | XP_026698469.1 |
| *Tyto alba alba* | XP_032865922.1 |
| *Dryobates pubescens* | XP_009899066.1 |
| *Leptosomus discolor* | XP_009947813.1 |
| *Accipiter nisus* | A0A8B9NMV5 |
| *Haliaeetus leucocephalus* | XP_010569558.1 |
| *Aquila chrysaetos chrysaetos* | XP_029875565.1 |
| *Nipponia nippon* | XP_009474527.1 |
| *Egretta garzetta* | XP_009647100.1 |
| *Pygoscelis adeliae* | XP_009323755.1 |
| *Aptenodytes forsteri* | XP_009275160.1 |
| *Limosa lapponica baueri* | A0A2I0UFW3 |
| *Calidris pugnax* | XP_014815788.1 |
| *Charadrius vociferus* | XP_009883023.1 |
| *Columba livia* | XP_005512804.1 |
| *Patagioenas fasciata monilis* | A0A1V4JBR9 |

| | |
|---|---|
| *Calypte anna* | XP_030316512.1 |
| *Cuculus canorus* | XP_009563852.1 |
| *Chlamydotis macqueenii* | XP_010114331.1 |
| *Anser cygnoides domesticus* | XP_013039377.1 |
| *Anser brachyrhynchus* | A0A8B9BYQ7 |
| *Cygnus olor* | XP_040388553.1 |
| *Cygnus atratus* | XP_035398131.1 |
| *Oxyura jamaicensis* | XP_035179769.1 |
| *Aythya fuligula* | XP_032037425.1 |
| *Anas platyrhynchos* | XP_027303973.2 |
| *Anas zonorhyncha* | A0A8B9VTS0 |
| *Cairina moschata domestica* | A0A8C3GFD3 |
| *Gallus gallus* | XP_015128429.2 |
| *Pavo cristatus* | A0A8C9FX21 |
| *Chrysolophus pictus* | A0A8C3LT74 |
| *Phasianus colchicus* | XP_031455223.1 |
| *Coturnix japonica* | XP_015741600.1 |
| *Numida meleagris* | XP_021265136.1 |
| *Apteryx owenii* | A0A8B9Q6I5 |
| *Apteryx mantelli mantelli* | XP_013807183.1 |
| *Dromaius novaehollandiae* | A0A8C4KFF6 |
| *Nothoprocta perdicaria* | XP_025907793.1 |
| *Struthio camelus australis* | XP_009667525.1 |
| *Alligator sinensis* | XP_006021289.1 |
| *Alligator mississippiensis* | XP_006272706.2 |
| *Gavialis gangeticus* | XP_019379795.1 |
| *Crocodylus porosus* | XP_019384784.1 |
| *Mauremys reevesii* | XP_039340216.1 |
| *Chelonoidis abingdonii* | XP_032619293.1 |
| *Trachemys scripta elegans* | XP_034611793.1 |
| *Dermochelys coriacea* | XP_038233949.1 |
| *Chelonia mydas* | XP_037757098.1 |
| *Chelydra serpentina* | A0A8C3RKC6 |
| *Pelodiscus sinensis* | XP_025035672.1 |
| *Pelusios castaneus* | A0A8C8SL48 |
| *Pseudonaja textilis* | XP_026569996.1 |
| *Notechis scutatus* | XP_026530732.1 |
| *Laticauda laticaudata* | A0A8C5RWC4 |
| *Naja naja* | A0A8C6Y605 |
| *Thamnophis sirtalis* | XP_013927292.1 |
| *Thamnophis elegans* | XP_032082409.1 |
| *Pantherophis guttatus* | XP_034290278.1 |
| *Crotalus tigris* | XP_039214321.1 |
| *Protobothrops mucrosquamatus* | XP_015680771.1 |
| *Python bivittatus* | XP_007439333.1 |
| *Pogona vitticeps* | XP_020644810.1 |
| *Anolis carolinensis* | XP_003218949.1 |
| *Varanus komodoensis* | A0A8D2J768 |

| | |
|---|---|
| *Zootoca vivipara* | XP_034970802.1 |
| *Lacerta agilis* | XP_033003220.1 |
| *Podarcis muralis* | XP_028583412.1 |
| *Salvator merianae* | A0A8D0EAZ8 |
| *Nanorana parkeri* | XP_018410344.1 |
| *Rana temporaria* | XP_040194146.1 |
| *Bufo bufo* | XP_040279629.1 |
| *Leptobrachium leishanense* | A0A8C5LY47 |
| *Xenopus tropicalis* | XP_002933671.2 |
| *Xenopus laevis* | A0A1L8HDF6 |
| *Microcaecilia unicolor* | XP_030058147.1 |
| *Geotrypetes seraphini* | XP_033806482.1 |
| *Rhinatrema bivittatum* | XP_029459197.1 |
| *Latimeria chalumnae* | XP_005997902.1 |
| *Nothobranchius furzeri* | A0A1A7ZVK7 |
| *Nothobranchius kadleci* | A0A1A8BXL8 |
| *Nothobranchius kuhntae* | A0A1A8J7T9 |
| *Nothobranchius rachovii* | A0A1A8NKB8 |
| *Nothobranchius pienaari* | A0A1A8L029 |
| *Nothobranchius korthausae* | A0A1A8EVJ2 |
| *Iconisemion striatum* | A0A1A7XIN0 |
| *Austrofundulus limnaeus* | A0A2I4ALT0 |
| *Kryptolebias marmoratus* | A0A3Q3B4A6 |
| *Nematolebias whitei* | XP_037546173.1 |
| *Xiphophorus couchianus* | XP_027870861.1 |
| *Xiphophorus maculatus* | XP_023188450.1 |
| *Xiphophorus hellerii* | XP_032417449.1 |
| *Poecilia formosa* | XP_007560217.1 |
| *Poecilia reticulata* | XP_008402733.1 |
| *Poecilia mexicana* | XP_014837042.1 |
| *Cyprinodon variegatus* | A0A3Q2C9M6 |
| *Fundulus heteroclitus* | XP_021178210.2 |
| *Oryzias latipes* | A0A3P9JNC1 |
| *Maylandia zebra* | XP_012772475.1 |
| *Astatotilapia calliptera* | XP_026020005.1 |
| *Pundamilia nyererei* | A0A3B4GTK5 |
| *Haplochromis burtoni* | XP_014195887.1 |
| *Neolamprologus brichardi* | XP_006780481.1 |
| *Oreochromis aureus* | A0A668S393 |
| *Oreochromis niloticus* | XP_005466862.1 |
| *Amphiprion ocellaris* | XP_023124169.1 |
| *Amphiprion percula* | A0A3P8TF53 |
| *Acanthochromis polyacanthus* | XP_022055688.1 |
| *Stegastes partitus* | A0A3B5ARI2 |
| *Salarias fasciatus* | XP_029960019.1 |
| *Gouania willdenowi* | XP_028298833.1 |
| *Parambassis ranga* | A0A6P7IBE4 |
| *Anabas testudineus* | XP_026233352.1 |

| | |
|---|---|
| *Betta splendens* | XP_028999450.1 |
| *Channa argus* | A0A6G1P915 |
| *Monopterus albus* | A0A3Q3JHU8 |
| *Scophthalmus maximus* | A0A2U9C452 |
| *Cynoglossus semilaevis* | A0A3P8V8K1 |
| *Lates calcarifer* | XP_018539179.1 |
| *Seriola dumerili* | A0A3B4TA94 |
| *Seriola lalandi dorsalis* | XP_023257449.1 |
| *Takifugu flavidus* | A0A5C6MT87 |
| *Sparus aurata* | XP_030269762.1 |
| *Larimichthys crocea* | XP_019133650.2 |
| *Dicentrarchus labrax* | A0A8C4EQJ0 |
| *Labrus bergylta* | A0A3Q3GQ68 |
| *Sander lucioperca* | XP_031162092.1 |
| *Perca flavescens* | XP_028438658.1 |
| *Etheostoma spectabile* | XP_032368750.1 |
| *Notothenia coriiceps* | XP_010778483.1 |
| *Gymnodraco acuticeps* | A0A6P8VKT0 |
| *Cyclopterus lumpus* | A0A8C2YVD5 |
| *Neogobius melanostomus* | A0A8C6U0K9 |
| *Hippocampus comes* | A0A3Q2ZBW1 |
| *Myripristis murdjan* | XP_029904469.1 |
| *Gadus morhua* | XP_030232503.1 |
| *Oncorhynchus kisutch* | A0A8C7K036 |
| *Salmo salar* | XP_014004951.1 |
| *Salmo trutta* | XP_029603501.1 |
| *Esox lucius* | XP_034145642.1 |
| *Cyprinus carpio* | XP_018940548.1 |
| *Carassius auratus* | A0A6P6J573 |
| *Danio rerio* | NP_001124243.1 |
| *Sinocyclocheilus anshuiensis* | XP_016345316.1 |
| *Sinocyclocheilus rhinocerous* | A0A673M732 |
| *Triplophysa tibetana* | A0A5A9NCK4 |
| *Ictalurus punctatus* | XP_017314736.1 |
| *Pangasianodon hypophthalmus* | XP_026803596.2 |
| *Tachysurus fulvidraco* | XP_027024345.1 |
| *Bagarius yarrelli* | A0A556U079 |
| *Astyanax mexicanus* | XP_007259435.2 |
| *Pygocentrus nattereri* | XP_017550134.1 |
| *Chanos chanos* | A0A6J2WFI7 |
| *Clupea harengus* | XP_031414946.1 |
| *Denticeps clupeoides* | XP_028835219.1 |
| *Lepisosteus oculatus* | XP_015219318.1 |
| *Erpetoichthys calabaricus* | XP_028655654.1 |
| *Amblyraja radiata* | XP_032888650.1 |
| *Rhincodon typus* | XP_020385617.1 |
| *Petromyzon marinus* | XP_032832891.1 |

| **Suppl. Table 2** - CDKL5 pathogenic coding variants (ClinVar database; Jul 27, 2024) | | | | | |
|---|---|---|---|---|---|
| **Name** | **Accession** | **VariationID** | **AlleleID(s)** | **dbSNP ID** | **Germline classification** |
| NM_001323289.2(CDKL5):c.53T>A (p.Val18Asp) | VCV000659752 | 659752 | 649896 | rs1602230515 | Pathogenic |
| NM_001323289.2(CDKL5):c.58G>C (p.Gly20Arg) | VCV000143828 | 143828 | 153560 | rs267608418 | Pathogenic |
| NM_001323289.2(CDKL5):c.59G>T (p.Gly20Val) | VCV000918032 | 918032 | 906351 | rs786204962 | Pathogenic |
| NM_001323289.2(CDKL5):c.61G>A (p.Glu21Lys) | VCV002951612 | 2951612 | 3114460 | | Pathogenic |
| NM_001323289.2(CDKL5):c.73G>A (p.Gly25Arg) | VCV000156659 | 156659 | 166513 | rs587783130 | Pathogenic |
| NM_001323289.2(CDKL5):c.74G>T (p.Gly25Val) | VCV002942811 | 2942811 | 3097397 | | Pathogenic |
| NM_001323289.2(CDKL5):c.80T>C (p.Val27Ala) | VCV001803084 | 1803084 | 1860113 | | Pathogenic/Likely pathogenic |
| NM_001323289.2(CDKL5):c.91A>G (p.Arg31Gly) | VCV000189599 | 189599 | 187580 | rs786204991 | Pathogenic |
| NM_001323289.2(CDKL5):c.119C>T (p.Ala40Val) | VCV000011502 | 11502 | 26541 | rs122460159 | Pathogenic |
| NM_001323289.2(CDKL5):c.125A>G (p.Lys42Arg) | VCV000143772 | 143772 | 153504 | rs267608429 | Pathogenic |
| NM_001323289.2(CDKL5):c.194G>C (p.Arg65Pro) | VCV001072615 | 1072615 | 1065250 | rs267608436 | Pathogenic |
| NM_001323289.2(CDKL5):c.211A>G (p.Asn71Asp) | VCV000156591 | 156591 | 166445 | rs587783072 | Pathogenic |
| NM_001323289.2(CDKL5):c.215T>A (p.Ile72Asn) | VCV000143797 | 143797 | 153529 | rs62641235 | Pathogenic |
| NM_001323289.2(CDKL5):c.215T>C (p.Ile72Thr) | VCV000011503 | 11503 | 26542 | rs62641235 | Pathogenic |
| NM_001323289.2(CDKL5):c.217G>A (p.Val73Met) | VCV001052408 | 1052408 | 1052361 | rs2147139605 | Pathogenic |
| NM_001323289.2(CDKL5):c.353A>G (p.Gln118Arg) | VCV000217875 | 217875 | 214546 | rs863225290 | Pathogenic |
| NM_001323289.2(CDKL5):c.356T>G (p.Leu119Arg) | VCV000156597 | 156597 | 166451 | rs587783078 | Pathogenic |
| NM_001323289.2(CDKL5):c.364G>A (p.Ala122Thr) | VCV000803716 | 803716 | 792186 | rs1602271692 | Pathogenic |
| NM_001323289.2(CDKL5):c.380A>G (p.His127Arg) | VCV000143818 | 143818 | 153550 | rs267608468 | Pathogenic |
| NM_001323289.2(CDKL5):c.401G>C (p.Arg134Pro) | VCV002578405 | 2578405 | 2743339 | | Pathogenic |
| NM_001323289.2(CDKL5):c.412C>G (p.Pro138Ala) | VCV001038592 | 1038592 | 1035436 | rs1925492354 | Pathogenic |
| NM_001323289.2(CDKL5):c.413C>T (p.Pro138Leu) | VCV000156600 | 156600 | 166454 | rs587783081 | Pathogenic |
| NM_001323289.2(CDKL5):c.416A>C (p.Glu139Ala) | VCV002942583 | 2942583 | 3097169 | | Pathogenic |
| NM_001323289.2(CDKL5):c.419A>G (p.Asn140Ser) | VCV000392860 | 392860 | 378016 | rs1057524663 | Pathogenic |
| NM_001323289.2(CDKL5):c.424T>G (p.Leu142Val) | VCV001037382 | 1037382 | 1035437 | rs1925493252 | Pathogenic |
| NM_001323289.2(CDKL5):c.449A>G (p.Lys150Arg) | VCV000156602 | 156602 | 166456 | rs587783083 | Pathogenic |
| NM_001323289.2(CDKL5):c.455G>A (p.Cys152Tyr) | VCV000495236 | 495236 | 486784 | rs122460157 | Pathogenic/Likely pathogenic |
| NM_001323289.2(CDKL5):c.458A>G (p.Asp153Gly) | VCV000189590 | 189590 | 187595 | rs786204985 | Pathogenic |
| NM_001323289.2(CDKL5):c.463G>A (p.Gly155Ser) | VCV002126431 | 2126431 | 2181731 | | Pathogenic |
| NM_001323289.2(CDKL5):c.514G>A (p.Val172Ile) | VCV000208653 | 208653 | 205338 | rs797044858 | Pathogenic |
| NM_001323289.2(CDKL5):c.515T>G (p.Val172Gly) | VCV002950640 | 2950640 | 3111610 | | Pathogenic |
| NM_001323289.2(CDKL5):c.523A>G (p.Arg175Gly) | VCV000983042 | 983042 | 971195 | rs1925574557 | Pathogenic |
| NM_001323289.2(CDKL5):c.525A>T (p.Arg175Ser) | VCV000011497 | 11497 | 26536 | rs61749700 | Pathogenic |
| NM_001323289.2(CDKL5):c.526T>G (p.Trp176Gly) | VCV000189594 | 189594 | 187600 | rs587783084 | Pathogenic |
| NM_001323289.2(CDKL5):c.528G>T (p.Trp176Cys) | VCV000189595 | 189595 | 187601 | rs786204989 | Pathogenic |
| NM_001323289.2(CDKL5):c.532C>G (p.Arg178Gly) | VCV001071237 | 1071237 | 1065253 | rs267608493 | Pathogenic |
| NM_001323289.2(CDKL5):c.532C>T (p.Arg178Trp) | VCV000143823 | 143823 | 153555 | rs267608493 | Pathogenic |
| NM_001323289.2(CDKL5):c.533G>T (p.Arg178Leu) | VCV002159555 | 2159555 | 1890561 | | Pathogenic |
| NM_001323289.2(CDKL5):c.533G>A (p.Arg178Gln) | VCV000094113 | 94113 | 100013 | rs267606715 | Pathogenic |
| NM_001323289.2(CDKL5):c.533G>C (p.Arg178Pro) | VCV000018450 | 18450 | 33489 | rs267606715 | Pathogenic |
| NM_001323289.2(CDKL5):c.539C>T (p.Pro180Leu) | VCV000143824 | 143824 | 153556 | rs61749704 | Pathogenic |
| NM_001323289.2(CDKL5):c.542A>C (p.Glu181Ala) | VCV000156604 | 156604 | 166458 | rs587783085 | Pathogenic |
| NM_001323289.2(CDKL5):c.545T>C (p.Leu182Pro) | VCV001700386 | 1700386 | 1692779 | rs2147145603 | Pathogenic |
| NM_001323289.2(CDKL5):c.577G>A (p.Asp193Asn) | VCV000156605 | 156605 | 166459 | rs587783086 | Pathogenic |
| NM_001323289.2(CDKL5):c.578A>C (p.Asp193Ala) | VCV001432338 | 1432338 | 1484449 | rs267608500 | Pathogenic |
| NM_001323289.2(CDKL5):c.583T>G (p.Trp195Gly) | VCV000567323 | 567323 | 572373 | rs1569215594 | Pathogenic |
| NM_001323289.2(CDKL5):c.587C>T (p.Ser196Leu) | VCV000143827 | 143827 | 153559 | rs267608501 | Pathogenic/Likely pathogenic |
| NM_001323289.2(CDKL5):c.593G>A (p.Gly198Asp) | VCV001006298 | 1006298 | 999706 | rs1925696959 | Pathogenic |
| NM_001323289.2(CDKL5):c.604G>T (p.Gly202Trp) | VCV001422104 | 1422104 | 1337718 | rs2147148032 | Pathogenic |
| NM_001323289.2(CDKL5):c.609G>C (p.Glu203Asp) | VCV002925658 | 2925658 | 3082653 | | Pathogenic |
| NM_001323289.2(CDKL5):c.620G>T (p.Gly207Val) | VCV001685608 | 1685608 | 1677624 | rs2147148055 | Pathogenic |
| NM_001323289.2(CDKL5):c.638G>A (p.Gly213Glu) | VCV000577127 | 577127 | 572376 | rs1569215629 | Pathogenic |
| NM_001323289.2(CDKL5):c.659T>C (p.Leu220Pro) | VCV000143830 | 143830 | 153562 | rs267608511 | Pathogenic |
| NM_001323289.2(CDKL5):c.812T>C (p.Leu271Pro) | VCV002138479 | 2138479 | 1868633 | | Pathogenic |
| NM_001323289.2(CDKL5):c.830T>C (p.Leu277Ser) | VCV001410163 | 1410163 | 1513471 | rs2147156094 | Pathogenic |
| NM_001323289.2(CDKL5):c.854G>A (p.Arg285Lys) | VCV000422272 | 422272 | 411215 | rs1064795672 | Pathogenic/Likely pathogenic |
| NM_001323289.2(CDKL5):c.863C>T (p.Thr288Ile) | VCV000011504 | 11504 | 26543 | rs267606713 | Pathogenic |
| NM_001323289.2(CDKL5):c.2152G>A (p.Val718Met) | VCV000143796 | 143796 | 153528 | rs267608653 | Pathogenic |
| NM_001323289.2(CDKL5):c.100G>T (p.Glu34Ter) | VCV002138478 | 2138478 | 1868631 | | Pathogenic |
| NM_001323289.2(CDKL5):c.154G>T (p.Glu52Ter) | VCV000861338 | 861338 | 849853 | rs1925261282 | Pathogenic |
| NM_001323289.2(CDKL5):c.163G>T (p.Glu55Ter) | VCV002944977 | 2944977 | 3107933 | | Pathogenic |
| NM_001323289.2(CDKL5):c.173T>A (p.Leu58Ter) | VCV000226417 | 226417 | 228220 | rs875989950 | Pathogenic |
| NM_001323289.2(CDKL5):c.175C>T (p.Arg59Ter) | VCV000143783 | 143783 | 153515 | rs62653623 | Pathogenic |
| NM_001323289.2(CDKL5):c.205C>T (p.Gln69Ter) | VCV001785194 | 1785194 | 1839991 | | Pathogenic |
| NM_001323289.2(CDKL5):c.220G>T (p.Glu74Ter) | VCV000156592 | 156592 | 166446 | rs587783073 | Pathogenic |
| NM_001323289.2(CDKL5):c.250A>T (p.Lys84Ter) | VCV000156593 | 156593 | 166447 | rs587783074 | Pathogenic |
| NM_001323289.2(CDKL5):c.258C>A (p.Tyr86Ter) | VCV002946090 | 2946090 | 3098083 | | Pathogenic |
| NM_001323289.2(CDKL5):c.258C>G (p.Tyr86Ter) | VCV000426143 | 426143 | 415757 | rs1085307470 | Pathogenic |
| NM_001323289.2(CDKL5):c.280A>T (p.Lys94Ter) | VCV001685606 | 1685606 | 1677622 | rs2147139684 | Pathogenic |
| NM_001323289.2(CDKL5):c.292G>T (p.Glu98Ter) | VCV001193529 | 1193529 | 1185760 | rs2147142586 | Pathogenic |
| NM_001323289.2(CDKL5):c.351T>A (p.Tyr117Ter) | VCV000156596 | 156596 | 166450 | rs587783077 | Pathogenic |

| | | | | | |
|---|---|---|---|---|---|
| NM_001323289.2(CDKL5):c.352C>T (p.Gln118Ter) | VCV000143817 | 143817 | 153549 | rs267608453 | Pathogenic |
| NM_001323289.2(CDKL5):c.400C>T (p.Arg134Ter) | VCV000143820 | 143820 | 153552 | rs267608472 | Pathogenic |
| NM_001323289.2(CDKL5):c.425T>G (p.Leu142Ter) | VCV000803718 | 803718 | 792188 | rs267608477 | Pathogenic |
| NM_001323289.2(CDKL5):c.425T>A (p.Leu142Ter) | VCV000143821 | 143821 | 153553 | rs267608477 | Pathogenic |
| NM_001323289.2(CDKL5):c.508G>T (p.Glu170Ter) | VCV000533384 | 533384 | 535118 | rs1555950066 | Pathogenic |
| NM_001323289.2(CDKL5):c.513C>G (p.Tyr171Ter) | VCV001072862 | 1072862 | 1065251 | rs267608490 | Pathogenic |
| NM_001323289.2(CDKL5):c.513C>A (p.Tyr171Ter) | VCV000143822 | 143822 | 153554 | rs267608490 | Pathogenic |
| NM_001323289.2(CDKL5):c.527G>A (p.Trp176Ter) | VCV001070360 | 1070360 | 1065252 | rs2147145565 | Pathogenic/Likely pathogenic |
| NM_001323289.2(CDKL5):c.585G>A (p.Trp195Ter) | VCV001456717 | 1456717 | 1454819 | rs2147147996 | Pathogenic |
| NM_001323289.2(CDKL5):c.607G>T (p.Glu203Ter) | VCV000143829 | 143829 | 153561 | rs267608505 | Pathogenic |
| NM_001323289.2(CDKL5):c.622C>T (p.Gln208Ter) | VCV000158186 | 158186 | 170101 | rs587783405 | Pathogenic |
| NM_001323289.2(CDKL5):c.670C>T (p.Gln224Ter) | VCV000916584 | 916584 | 904944 | rs1925701271 | Pathogenic |
| NM_001323289.2(CDKL5):c.697G>T (p.Glu233Ter) | VCV000620492 | 620492 | 611937 | rs1569215664 | Pathogenic |
| NM_001323289.2(CDKL5):c.700C>T (p.Gln234Ter) | VCV000156608 | 156608 | 166462 | rs587783089 | Pathogenic |
| NM_001323289.2(CDKL5):c.766C>T (p.Gln256Ter) | VCV000503610 | 503610 | 495894 | rs1555951142 | Pathogenic |
| NM_001323289.2(CDKL5):c.786C>A (p.Tyr262Ter) | VCV000489166 | 489166 | 482246 | rs1555951146 | Pathogenic |
| NM_001323289.2(CDKL5):c.858C>A (p.Tyr286Ter) | VCV000419625 | 419625 | 411216 | rs766511365 | Pathogenic |
| NM_001323289.2(CDKL5):c.868C>T (p.Gln290Ter) | VCV000560971 | 560971 | 552253 | rs1569218019 | Pathogenic |
| NM_001323289.2(CDKL5):c.934A>T (p.Lys312Ter) | VCV002035673 | 2035673 | 2097457 | | Pathogenic |
| NM_001323289.2(CDKL5):c.968T>A (p.Leu323Ter) | VCV002134748 | 2134748 | 2192305 | | Pathogenic |
| NM_001323289.2(CDKL5):c.1006C>T (p.Gln336Ter) | VCV000373086 | 373086 | 360565 | rs1057518203 | Pathogenic |
| NM_001323289.2(CDKL5):c.1039C>T (p.Gln347Ter) | VCV000143767 | 143767 | 153499 | rs267608561 | Pathogenic |
| NM_001323289.2(CDKL5):c.1090G>T (p.Glu364Ter) | VCV000189553 | 189553 | 187610 | rs786204966 | Pathogenic |
| NM_001323289.2(CDKL5):c.1152C>G (p.Tyr384Ter) | VCV000194049 | 194049 | 191212 | rs794727064 | Pathogenic |
| NM_001323289.2(CDKL5):c.1165C>T (p.Gln389Ter) | VCV000590188 | 590188 | 580914 | rs1569219346 | Pathogenic |
| NM_001323289.2(CDKL5):c.1238C>G (p.Ser413Ter) | VCV000143771 | 143771 | 153503 | rs267608618 | Pathogenic |
| NM_001323289.2(CDKL5):c.1246G>T (p.Glu416Ter) | VCV000464808 | 464808 | 470650 | rs1555951991 | Pathogenic |
| NM_001323289.2(CDKL5):c.1321C>T (p.Gln441Ter) | VCV002123296 | 2123296 | 2177195 | | Pathogenic |
| NM_001323289.2(CDKL5):c.1324C>T (p.Gln442Ter) | VCV000156682 | 156682 | 166536 | rs587783151 | Pathogenic |
| NM_001323289.2(CDKL5):c.1345G>T (p.Glu449Ter) | VCV000434663 | 434663 | 430758 | rs1555952015 | Pathogenic |
| NM_001323289.2(CDKL5):c.1354C>T (p.Gln452Ter) | VCV000817555 | 817555 | 806179 | rs1602286285 | Pathogenic |
| NM_001323289.2(CDKL5):c.1375C>T (p.Gln459Ter) | VCV000189556 | 189556 | 187614 | rs786204969 | Pathogenic |
| NM_001323289.2(CDKL5):c.1486A>T (p.Lys496Ter) | VCV000870174 | 870174 | 858373 | rs1926283887 | Pathogenic |
| NM_001323289.2(CDKL5):c.1519C>T (p.Gln507Ter) | VCV000464809 | 464809 | 471431 | rs1555952052 | Pathogenic |
| NM_001323289.2(CDKL5):c.1548C>A (p.Tyr516Ter) | VCV001685611 | 1685611 | 1677627 | rs751789670 | Pathogenic |
| NM_001323289.2(CDKL5):c.1648C>T (p.Arg550Ter) | VCV000143780 | 143780 | 153512 | rs267608643 | Pathogenic |
| NM_001323289.2(CDKL5):c.1670C>G (p.Ser557Ter) | VCV000156676 | 156676 | 166530 | rs587783145 | Pathogenic |
| NM_001323289.2(CDKL5):c.1675C>T (p.Arg559Ter) | VCV000143781 | 143781 | 153513 | rs267608395 | Pathogenic/Likely pathogenic |
| NM_001323289.2(CDKL5):c.1708G>T (p.Glu570Ter) | VCV000143782 | 143782 | 153514 | rs267608644 | Pathogenic |
| NM_001323289.2(CDKL5):c.1813C>T (p.Gln605Ter) | VCV000976039 | 976039 | 964590 | rs1926301954 | Pathogenic |
| NM_001323289.2(CDKL5):c.1816C>T (p.Gln606Ter) | VCV000803723 | 803723 | 792193 | rs1602286899 | Pathogenic |
| NM_001323289.2(CDKL5):c.1897C>T (p.Gln633Ter) | VCV000217357 | 217357 | 196443 | rs863225065 | Pathogenic |
| NM_001323289.2(CDKL5):c.1927C>T (p.Gln643Ter) | VCV001070908 | 1070908 | 1065257 | rs2147161353 | Pathogenic |
| NM_001323289.2(CDKL5):c.1954C>T (p.Gln652Ter) | VCV000143790 | 143790 | 153522 | rs267608647 | Pathogenic |
| NM_001323289.2(CDKL5):c.2095G>T (p.Glu699Ter) | VCV000280475 | 280475 | 265145 | rs886041673 | Pathogenic |
| NM_001323289.2(CDKL5):c.2277G>A (p.Trp759Ter) | VCV002941715 | 2941715 | 3107246 | | Pathogenic |
| NM_001323289.2(CDKL5):c.2305G>T (p.Glu769Ter) | VCV000280719 | 280719 | 264971 | rs886041872 | Pathogenic |
| NM_001323289.2(CDKL5):c.2345C>G (p.Ser782Ter) | VCV000662018 | 662018 | 649906 | rs1555954074 | Pathogenic |
| NM_001323289.2(CDKL5):c.2345C>A (p.Ser782Ter) | VCV000464812 | 464812 | 471434 | rs1555954074 | Pathogenic/Likely pathogenic |
| NM_001323289.2(CDKL5):c.2371C>T (p.Gln791Ter) | VCV001361871 | 1361871 | 1376497 | rs1926838663 | Pathogenic |
| NM_001323289.2(CDKL5):c.2413C>T (p.Gln805Ter) | VCV000143804 | 143804 | 153536 | rs267608659 | Pathogenic/Likely pathogenic |
| NM_001323289.2(CDKL5):c.2480C>G (p.Ser827Ter) | VCV000803724 | 803724 | 792194 | rs1602298653 | Pathogenic |
| NM_001323289.2(CDKL5):c.2494C>T (p.Gln832Ter) | VCV000189573 | 189573 | 187625 | rs17857094 | Pathogenic |
| NM_001323289.2(CDKL5):c.2578C>T (p.Gln860Ter) | VCV000916583 | 916583 | 904946 | rs1927139054 | Pathogenic |
| NM_001323289.2(CDKL5):c.2593C>T (p.Gln865Ter) | VCV000143808 | 143808 | 153540 | rs267608663 | Pathogenic |
| NM_001323289.2(CDKL5):c.2596C>T (p.Gln866Ter) | VCV000156691 | 156691 | 166545 | rs587783158 | Pathogenic |
| NM_001323289.2(CDKL5):c.2641C>T (p.Gln881Ter) | VCV000375523 | 375523 | 362353 | rs1057519541 | Pathogenic |
| NM_001323289.2(CDKL5):c.2671C>T (p.Gln891Ter) | VCV000156692 | 156692 | 166546 | rs587783159 | Pathogenic |
| NM_001323289.2(CDKL5):c.2704C>T (p.Gln902Ter) | VCV000189583 | 189583 | 187628 | rs786204981 | Pathogenic |
| NM_001323289.2(CDKL5):c.2716C>T (p.Gln906Ter) | VCV000217874 | 217874 | 214547 | rs863225289 | Pathogenic |
| NM_001323289.2(CDKL5):c.2785C>T (p.Gln929Ter) | VCV000648838 | 648838 | 649910 | rs1602300816 | Pathogenic |
| NM_001323289.2(CDKL5):c.2842C>T (p.Arg948Ter) | VCV000489299 | 489299 | 482247 | rs1555955296 | Pathogenic |
| NM_001323289.2(CDKL5):c.39del (p.Phe13fs) | VCV000143819 | 143819 | 153551 | rs267608415 | Pathogenic |
| NM_001323289.2(CDKL5):c.46del (p.Val18fs) | VCV002942111 | 2942111 | 3104456 | | Pathogenic |
| NM_001323289.2(CDKL5):c.155_156del (p.Glu52fs) | VCV001338216 | 1338216 | 1329225 | rs2147139532 | Pathogenic |
| NM_001323289.2(CDKL5):c.163_166del (p.Glu55fs) | VCV000143779 | 143779 | 153511 | rs267608433 | Pathogenic/Likely pathogenic |
| NM_001323289.2(CDKL5):c.183del (p.Met63fs) | VCV000143785 | 143785 | 153517 | rs62643608 | Pathogenic |
| NM_001323289.2(CDKL5):c.200_201del (p.Leu67fs) | VCV000533386 | 533386 | 534765 | rs1555949011 | Pathogenic |
| NM_001323289.2(CDKL5):c.207_213del (p.Glu70fs) | VCV000189569 | 189569 | 187584 | rs786204977 | Pathogenic |
| NM_001323289.2(CDKL5):c.212del (p.Asn71fs) | VCV000286056 | 286056 | 270293 | rs886043296 | Pathogenic |
| NM_001323289.2(CDKL5):c.229_232del (p.Glu77fs) | VCV000143799 | 143799 | 153531 | rs267608441 | Pathogenic |
| NM_001323289.2(CDKL5):c.244del (p.Arg82fs) | VCV000464813 | 464813 | 471428 | rs1555949041 | Pathogenic |
| NM_001323289.2(CDKL5):c.248del (p.Gly83fs) | VCV000156637 | 156637 | 166491 | rs587783109 | Pathogenic |

| | | | | | |
|---|---|---|---|---|---|
| NM_001323289.2(CDKL5):c.275_276insAA (p.Glu93fs) | VCV000189584 | 189584 | 187586 | rs786204982 | Pathogenic |
| NM_001323289.2(CDKL5):c.308del (p.Met103fs) | VCV001456413 | 1456413 | 1458352 | rs2147142599 | Pathogenic |
| NM_001323289.2(CDKL5):c.349_352del (p.Tyr117fs) | VCV001685607 | 1685607 | 1677623 | rs2147142645 | Pathogenic |
| NM_001323289.2(CDKL5):c.349dup (p.Tyr117fs) | VCV000870178 | 870178 | 858370 | rs1925420814 | Pathogenic |
| NM_001323289.2(CDKL5):c.354_361del (p.Leu119fs) | VCV000870351 | 870351 | 858516 | rs1925421198 | Pathogenic |
| NM_001323289.2(CDKL5):c.358_367del (p.Ile120fs) | VCV000987448 | 987448 | 975607 | rs1925421858 | Pathogenic |
| NM_001323289.2(CDKL5):c.372_385del (p.His124fs) | VCV002105592 | 2105592 | 2169944 | | Pathogenic |
| NM_001323289.2(CDKL5):c.383del (p.Lys128fs) | VCV000817775 | 817775 | 806176 | rs1602271715 | Pathogenic |
| NM_001323289.2(CDKL5):c.386del (p.Asn129fs) | VCV000803717 | 803717 | 792187 | rs1602271718 | Pathogenic |
| NM_001323289.2(CDKL5):c.427dup (p.Ile143fs) | VCV000432570 | 432570 | 426444 | rs1555949752 | Pathogenic |
| NM_001323289.2(CDKL5):c.450del (p.Lys150fs) | VCV001407573 | 1407573 | 1506454 | rs2147144049 | Pathogenic |
| NM_001323289.2(CDKL5):c.453del (p.Cys152fs) | VCV000488477 | 488477 | 481345 | rs1555949763 | Pathogenic |
| NM_001323289.2(CDKL5):c.495dup (p.Ala166fs) | VCV000870173 | 870173 | 858372 | rs1925573011 | Pathogenic |
| NM_001323289.2(CDKL5):c.506_507del (p.Thr169fs) | VCV000189592 | 189592 | 187598 | rs786204987 | Pathogenic |
| NM_001323289.2(CDKL5):c.510_511dup (p.Tyr171fs) | VCV000189593 | 189593 | 187599 | rs786204988 | Pathogenic |
| NM_001323289.2(CDKL5):c.549_552del (p.Leu184fs) | VCV000156639 | 156639 | 166493 | rs587783111 | Pathogenic |
| NM_001323289.2(CDKL5):c.549dup (p.Leu184fs) | VCV000143825 | 143825 | 153557 | rs267608497 | Pathogenic |
| NM_001323289.2(CDKL5):c.614_617dup (p.Asp206fs) | VCV000533389 | 533389 | 535119 | rs1555950465 | Pathogenic |
| NM_001323289.2(CDKL5):c.629del (p.Leu210fs) | VCV000524152 | 524152 | 514794 | rs1555950470 | Pathogenic |
| NM_001323289.2(CDKL5):c.630_631del (p.Leu210fs) | VCV001804081 | 1804081 | 1861106 | | Pathogenic |
| NM_001323289.2(CDKL5):c.663dup (p.Thr222fs) | VCV000817077 | 817077 | 806177 | rs1602276166 | Pathogenic |
| NM_001323289.2(CDKL5):c.660_664dup (p.Thr222fs) | VCV000189597 | 189597 | 187604 | rs786204990 | Pathogenic |
| NM_001323289.2(CDKL5):c.665del (p.Thr222fs) | VCV001723885 | 1723885 | 1781370 | | Pathogenic |
| NM_001323289.2(CDKL5):c.666del (p.Ile223fs) | VCV000587758 | 587758 | 580892 | rs1569215645 | Pathogenic |
| NM_001323289.2(CDKL5):c.692del (p.Pro231fs) | VCV002927402 | 2927402 | 3083085 | | Pathogenic |
| NM_001323289.2(CDKL5):c.693del (p.Ser232fs) | VCV000280587 | 280587 | 264970 | rs886041764 | Pathogenic |
| NM_001323289.2(CDKL5):c.712_713dup (p.Tyr239fs) | VCV002038957 | 2038957 | 2098150 | | Pathogenic |
| NM_001323289.2(CDKL5):c.713del (p.Phe238fs) | VCV000501058 | 501058 | 492482 | rs1555950486 | Pathogenic |
| NM_001323289.2(CDKL5):c.725del (p.Pro242fs) | VCV000533390 | 533390 | 535120 | rs1555950494 | Pathogenic |
| NM_001323289.2(CDKL5):c.747dup (p.Pro250fs) | VCV000226116 | 226116 | 227924 | rs875989880 | Pathogenic |
| NM_001323289.2(CDKL5):c.749del (p.Pro250fs) | VCV001456215 | 1456215 | 1458155 | rs2147153531 | Pathogenic |
| NM_001323289.2(CDKL5):c.751delinsTC (p.Ala251fs) | VCV000523629 | 523629 | 514279 | rs1555951141 | Pathogenic |
| NM_001323289.2(CDKL5):c.768dup (p.Ser257fs) | VCV000430365 | 430365 | 422445 | rs1131691926 | Pathogenic |
| NM_001323289.2(CDKL5):c.781dup (p.Arg261fs) | VCV002945065 | 2945065 | 3108021 | | Pathogenic |
| NM_001323289.2(CDKL5):c.801_802del (p.Asn267fs) | VCV000143833 | 143833 | 153565 | rs267608528 | Pathogenic |
| NM_001323289.2(CDKL5):c.808dup (p.Leu270fs) | VCV000156640 | 156640 | 166494 | rs587783112 | Pathogenic |
| NM_001323289.2(CDKL5):c.838_847del (p.Asp281fs) | VCV000143834 | 143834 | 153566 | rs61750250 | Pathogenic |
| NM_001323289.2(CDKL5):c.867dup (p.Gln290fs) | VCV000143836 | 143836 | 153568 | rs267608537 | Pathogenic |
| NM_001323289.2(CDKL5):c.878dup (p.Asn293fs) | VCV000970805 | 970805 | 959217 | rs1926080187 | Pathogenic |
| NM_001323289.2(CDKL5):c.877_881dup (p.His294fs) | VCV000845526 | 845526 | 849859 | rs1926080344 | Pathogenic |
| NM_001323289.2(CDKL5):c.884del (p.Pro295fs) | VCV000143837 | 143837 | 153569 | rs267608542 | Pathogenic |
| NM_001323289.2(CDKL5):c.898_899del (p.Gln300fs) | VCV000813762 | 813762 | 802048 | rs1602282699 | Pathogenic |
| NM_001323289.2(CDKL5):c.902_903dup (p.Leu302fs) | VCV000143838 | 143838 | 153570 | rs267608546 | Pathogenic |
| NM_001323289.2(CDKL5):c.906del (p.Leu303fs) | VCV001685609 | 1685609 | 1677625 | rs2147156191 | Pathogenic |
| NM_001323289.2(CDKL5):c.907_908insCC (p.Leu303fs) | VCV000817286 | 817286 | 806178 | rs1602282705 | Pathogenic |
| NM_001323289.2(CDKL5):c.942del (p.Lys314fs) | VCV000189600 | 189600 | 187606 | rs786204992 | Pathogenic |
| NM_001323289.2(CDKL5):c.964dup (p.Thr322fs) | VCV000143840 | 143840 | 153572 | rs267608552 | Pathogenic |
| NM_001323289.2(CDKL5):c.989_1010del (p.Gly330fs) | VCV000871042 | 871042 | 860819 | rs1926254082 | Pathogenic |
| NM_001323289.2(CDKL5):c.993_994dup (p.Ser332fs) | VCV001354281 | 1354281 | 1341399 | rs2147160115 | Pathogenic |
| NM_001323289.2(CDKL5):c.1007_1014del (p.Gln336fs) | VCV000418707 | 418707 | 411219 | rs1064793381 | Pathogenic |
| NM_001323289.2(CDKL5):c.1008del (p.Gln336fs) | VCV000988540 | 988540 | 976490 | rs1926255341 | Pathogenic |
| NM_001323289.2(CDKL5):c.1008_1029del (p.Ser337fs) | VCV000189551 | 189551 | 187608 | rs786204964 | Pathogenic |
| NM_001323289.2(CDKL5):c.1011dup (p.His338fs) | VCV001451469 | 1451469 | 1425871 | rs2147160162 | Pathogenic |
| NM_001323289.2(CDKL5):c.1053_1056dup (p.Leu353fs) | VCV001072660 | 1072660 | 1065255 | rs2147160237 | Pathogenic |
| NM_001323289.2(CDKL5):c.1066dup (p.Ala356fs) | VCV001350934 | 1350934 | 1492522 | rs2147160283 | Pathogenic |
| NM_001323289.2(CDKL5):c.1071del (p.Asp357fs) | VCV000189552 | 189552 | 187609 | rs786204965 | Pathogenic |
| NM_001323289.2(CDKL5):c.1082dup (p.Ala362fs) | VCV000143769 | 143769 | 153501 | rs267608566 | Pathogenic |
| NM_001323289.2(CDKL5):c.1079del (p.Leu360fs) | VCV000143768 | 143768 | 153500 | rs267608565 | Pathogenic |
| NM_001323289.2(CDKL5):c.1094dup (p.Ser365fs) | VCV000975929 | 975929 | 964589 | rs1926261579 | Pathogenic |
| NM_001323289.2(CDKL5):c.1103dup (p.Asn368fs) | VCV000981157 | 981157 | 969243 | rs1926262196 | Pathogenic |
| NM_001323289.2(CDKL5):c.1108_1109dup (p.Asn370fs) | VCV000287828 | 287828 | 272065 | rs886043742 | Pathogenic |
| NM_001323289.2(CDKL5):c.1111del (p.Ala372fs) | VCV000502250 | 502250 | 493674 | rs1555951958 | Pathogenic |
| NM_001323289.2(CDKL5):c.1166_1169del (p.Gln389fs) | VCV002099555 | 2099555 | 2147113 | | Pathogenic |
| NM_001323289.2(CDKL5):c.1180_1181dup (p.Ser394fs) | VCV002944760 | 2944760 | 3107717 | | Pathogenic |
| NM_001323289.2(CDKL5):c.1208del (p.Pro403fs) | VCV002033947 | 2033947 | 2097007 | | Pathogenic |
| NM_001323289.2(CDKL5):c.1211_1212dup (p.Leu405fs) | VCV000423029 | 423029 | 411222 | rs1555951981 | Pathogenic |
| NM_001323289.2(CDKL5):c.1247_1248del (p.Glu416fs) | VCV000189554 | 189554 | 187611 | rs786204967 | Pathogenic |
| NM_001323289.2(CDKL5):c.1287dup (p.Gly430fs) | VCV002921376 | 2921376 | 3088754 | | Pathogenic |
| NM_001323289.2(CDKL5):c.1311dup (p.Ser438fs) | VCV000143774 | 143774 | 153506 | rs267608623 | Pathogenic |
| NM_001323289.2(CDKL5):c.1330_1342del (p.Arg444fs) | VCV002953180 | 2953180 | 3114735 | | Pathogenic |
| NM_001323289.2(CDKL5):c.1341del (p.Phe447fs) | VCV000189555 | 189555 | 187613 | rs786204968 | Pathogenic |
| NM_001323289.2(CDKL5):c.1345_1363del (p.Glu449fs) | VCV000156641 | 156641 | 166495 | rs587783113 | Pathogenic |
| NM_001323289.2(CDKL5):c.1345_1346del (p.Glu449fs) | VCV000158177 | 158177 | 170104 | rs587783398 | Pathogenic |

| | | | | | |
|---|---|---|---|---|---|
| NM_001323289.2(CDKL5):c.1365_1366insA (p.Gly456fs) | VCV000290315 | 290315 | 274552 | rs886044424 | Pathogenic |
| NM_001323289.2(CDKL5):c.1417dup (p.Ile473fs) | VCV000189557 | 189557 | 187615 | rs786204970 | Pathogenic |
| NM_001323289.2(CDKL5):c.1419del (p.Gln475fs) | VCV001342902 | 1342902 | 1334541 | rs2147160726 | Pathogenic |
| NM_001323289.2(CDKL5):c.1431_1435dup (p.Ser479fs) | VCV000803720 | 803720 | 792190 | rs1602286391 | Pathogenic |
| NM_001323289.2(CDKL5):c.1432_1433insT (p.Arg478fs) | VCV000189558 | 189558 | 187616 | rs786204971 | Pathogenic |
| NM_001323289.2(CDKL5):c.1449_1452dup (p.Lys485fs) | VCV000375524 | 375524 | 362352 | rs1057519542 | Pathogenic |
| NM_001323289.2(CDKL5):c.1452del (p.Lys485fs) | VCV002944504 | 2944504 | 3097793 | | Pathogenic |
| NM_001323289.2(CDKL5):c.1471dup (p.Ala491fs) | VCV000810521 | 810521 | 798246 | rs1602286449 | Pathogenic |
| NM_001323289.2(CDKL5):c.1485dup (p.Lys496fs) | VCV000838071 | 838071 | 849861 | rs1926283743 | Pathogenic |
| NM_001323289.2(CDKL5):c.1546del (p.Tyr516fs) | VCV000503833 | 503833 | 495794 | rs1555952063 | Pathogenic |
| NM_001323289.2(CDKL5):c.1550del (p.Phe517fs) | VCV000189560 | 189560 | 187617 | rs786204972 | Pathogenic |
| NM_001323289.2(CDKL5):c.1553del (p.Pro518fs) | VCV000284540 | 284540 | 268777 | rs886042899 | Pathogenic |
| NM_001323289.2(CDKL5):c.1584dup (p.Ser529fs) | VCV001072905 | 1072905 | 1065256 | rs2147160886 | Pathogenic |
| NM_001323289.2(CDKL5):c.1612_1613del (p.Thr538fs) | VCV001335976 | 1335976 | 1326989 | rs2147160920 | Pathogenic |
| NM_001323289.2(CDKL5):c.1671dup (p.Arg558fs) | VCV000156643 | 156643 | 166497 | rs587783115 | Pathogenic/Likely pathogenic |
| NM_001323289.2(CDKL5):c.1684_1687del (p.Thr562fs) | VCV000434665 | 434665 | 430761 | rs1555952101 | Pathogenic |
| NM_001323289.2(CDKL5):c.1686del (p.Arg563fs) | VCV001422610 | 1422610 | 1412504 | rs2147161009 | Pathogenic |
| NM_001323289.2(CDKL5):c.1741del (p.His581fs) | VCV002951292 | 2951292 | 3114140 | | Pathogenic |
| NM_001323289.2(CDKL5):c.1742dup (p.His581fs) | VCV000803722 | 803722 | 792192 | rs1602286792 | Pathogenic |
| NM_001323289.2(CDKL5):c.1744dup (p.Ser582fs) | VCV001879752 | 1879752 | 1936578 | | Pathogenic |
| NM_001323289.2(CDKL5):c.1756_1759del (p.Ser586fs) | VCV000916588 | 916588 | 904945 | rs1926298368 | Pathogenic |
| NM_001323289.2(CDKL5):c.1755_1756del (p.Ser586fs) | VCV000856634 | 856634 | 849862 | rs1926298521 | Pathogenic |
| NM_001323289.2(CDKL5):c.1776_1777del (p.Ser593fs) | VCV000422734 | 422734 | 411224 | rs1064793376 | Pathogenic |
| NM_001323289.2(CDKL5):c.1777del (p.Ser593fs) | VCV000418699 | 418699 | 411225 | rs1064793376 | Pathogenic |
| NM_001323289.2(CDKL5):c.1784dup (p.Leu596fs) | VCV000189566 | 189566 | 187618 | rs786204974 | Pathogenic |
| NM_001323289.2(CDKL5):c.1790del (p.Gly597fs) | VCV002001461 | 2001461 | 2054662 | | Pathogenic |
| NM_001323289.2(CDKL5):c.1795dup (p.Thr599fs) | VCV000156644 | 156644 | 166498 | rs587783116 | Pathogenic |
| NM_001323289.2(CDKL5):c.1797dup (p.Ser600fs) | VCV000158181 | 158181 | 170107 | rs587783401 | Pathogenic |
| NM_001323289.2(CDKL5):c.1852_1853insT (p.Asp618fs) | VCV000870169 | 870169 | 858374 | rs1926303951 | Pathogenic |
| NM_001323289.2(CDKL5):c.1854del (p.Asp618fs) | VCV000189567 | 189567 | 187619 | rs786204975 | Pathogenic |
| NM_001323289.2(CDKL5):c.1883del (p.Ser628fs) | VCV000840609 | 840609 | 849863 | rs1926305134 | Pathogenic |
| NM_001323289.2(CDKL5):c.1885_1886dup (p.Leu629fs) | VCV000156645 | 156645 | 166499 | rs587783117 | Pathogenic |
| NM_001323289.2(CDKL5):c.1891_1916del (p.Ile631fs) | VCV000194048 | 194048 | 191211 | rs794727063 | Pathogenic |
| NM_001323289.2(CDKL5):c.1896del (p.Gln633fs) | VCV001400702 | 1400702 | 1463554 | rs2147161299 | Pathogenic |
| NM_001323289.2(CDKL5):c.1909del (p.Ala637fs) | VCV000156646 | 156646 | 166500 | rs587783118 | Pathogenic |
| NM_001323289.2(CDKL5):c.1921_1922del (p.Ser641fs) | VCV000560636 | 560636 | 551754 | rs1569219844 | Pathogenic |
| NM_001323289.2(CDKL5):c.1976_1977del (p.Val659fs) | VCV001072616 | 1072616 | 1065258 | rs2147164191 | Pathogenic |
| NM_001323289.2(CDKL5):c.1994_1997del (p.Lys665fs) | VCV002497678 | 2497678 | 2472761 | | Pathogenic |
| NM_001323289.2(CDKL5):c.1994_1995del (p.Lys665fs) | VCV001784021 | 1784021 | 1849580 | | Pathogenic |
| NM_001323289.2(CDKL5):c.2009_2012dup (p.Thr672fs) | VCV002026466 | 2026466 | 2088222 | | Pathogenic |
| NM_001323289.2(CDKL5):c.2016dup (p.Ser673fs) | VCV000143793 | 143793 | 153525 | rs267608648 | Pathogenic |
| NM_001323289.2(CDKL5):c.2016del (p.Ser673fs) | VCV000143792 | 143792 | 153524 | rs267608648 | Pathogenic |
| NM_001323289.2(CDKL5):c.2022del (p.Phe675fs) | VCV000408123 | 408123 | 404163 | rs1060501860 | Pathogenic |
| NM_001323289.2(CDKL5):c.2026del (p.His676fs) | VCV000947188 | 947188 | 929637 | rs1926450614 | Pathogenic |
| NM_001323289.2(CDKL5):c.2066del (p.Pro689fs) | VCV000143795 | 143795 | 153527 | rs267608651 | Pathogenic |
| NM_001323289.2(CDKL5):c.2091del (p.Glu699fs) | VCV002127982 | 2127982 | 2185859 | | Pathogenic |
| NM_001323289.2(CDKL5):c.2094del (p.Glu699fs) | VCV000992590 | 992590 | 980515 | rs1926473501 | Pathogenic |
| NM_001323289.2(CDKL5):c.2101dup (p.Arg701fs) | VCV001676144 | 1676144 | 1667525 | rs2147164710 | Pathogenic |
| NM_001323289.2(CDKL5):c.2105_2106del (p.His702fs) | VCV000189570 | 189570 | 187621 | rs786204978 | Pathogenic |
| NM_001323289.2(CDKL5):c.2142del (p.Tyr716fs) | VCV000576176 | 576176 | 573755 | rs1569220925 | Pathogenic |
| NM_001323289.2(CDKL5):c.2182del (p.His728fs) | VCV000870175 | 870175 | 858375 | rs1926616715 | Pathogenic |
| NM_001323289.2(CDKL5):c.2197_2204dup (p.Arg735fs) | VCV000857515 | 857515 | 849864 | rs1926617807 | Pathogenic |
| NM_001323289.2(CDKL5):c.2205_2206del (p.Arg735fs) | VCV000156648 | 156648 | 166502 | rs587783120 | Pathogenic |
| NM_001323289.2(CDKL5):c.2216del (p.Leu739fs) | VCV001787844 | 1787844 | 1847287 | | Pathogenic |
| NM_001323289.2(CDKL5):c.2227_2228del (p.Ser743fs) | VCV001451225 | 1451225 | 1429479 | rs2147167604 | Pathogenic |
| NM_001323289.2(CDKL5):c.2231_2265del (p.Ser744fs) | VCV000423577 | 423577 | 411229 | rs1064796503 | Pathogenic |
| NM_001323289.2(CDKL5):c.2247_2250dup (p.Lys751fs) | VCV002952563 | 2952563 | 3110177 | | Pathogenic |
| NM_001323289.2(CDKL5):c.2254dup (p.Arg752fs) | VCV000156649 | 156649 | 166503 | rs587783121 | Pathogenic |
| NM_001323289.2(CDKL5):c.2255_2259dup (p.Pro754fs) | VCV002921951 | 2921951 | 3085442 | | Pathogenic |
| NM_001323289.2(CDKL5):c.2256_2263del (p.Arg752fs) | VCV000969040 | 969040 | 959219 | rs1926620965 | Pathogenic |
| NM_001323289.2(CDKL5):c.2323_2326del (p.Glu775fs) | VCV000156650 | 156650 | 166504 | rs267608654 | Pathogenic |
| NM_001323289.2(CDKL5):c.2325_2326del (p.Lys776fs) | VCV000143800 | 143800 | 153532 | rs267608654 | Pathogenic |
| NM_001323289.2(CDKL5):c.2326_2327del (p.Lys776fs) | VCV000652635 | 652635 | 649905 | rs1602295727 | Pathogenic |
| NM_001323289.2(CDKL5):c.2343del (p.Arg781fs) | VCV000143801 | 143801 | 153533 | rs62643614 | Pathogenic |
| NM_001323289.2(CDKL5):c.2355_2358del (p.Lys786fs) | VCV000870179 | 870179 | 858376 | rs1926837956 | Pathogenic |
| NM_001323289.2(CDKL5):c.2359_2363del (p.Lys787fs) | VCV000437405 | 437405 | 430972 | rs1555954078 | Pathogenic |
| NM_001323289.2(CDKL5):c.2360_2363del (p.Lys787fs) | VCV000156651 | 156651 | 166505 | rs587783123 | Pathogenic |
| NM_001323289.2(CDKL5):c.2363_2367del (p.Lys788fs) | VCV000143802 | 143802 | 153534 | rs267608655 | Pathogenic |
| NM_001323289.2(CDKL5):c.2374dup (p.Thr792fs) | VCV000817613 | 817613 | 806180 | rs1602295779 | Pathogenic/Likely pathogenic |
| NM_001323289.2(CDKL5):c.2398del (p.Asp800fs) | VCV002945903 | 2945903 | 3101783 | | Pathogenic |
| NM_001323289.2(CDKL5):c.2432del (p.Ala811fs) | VCV000916505 | 916505 | 904726 | rs1927001903 | Pathogenic |
| NM_001323289.2(CDKL5):c.2452_2459del (p.Pro818fs) | VCV000548518 | 548518 | 539112 | rs1555954737 | Pathogenic |
| NM_001323289.2(CDKL5):c.2463del (p.Trp821fs) | VCV001376941 | 1376941 | 1394556 | rs2147175902 | Pathogenic |

| | | | | | |
|---|---|---|---|---|---|
| NM_001323289.2(CDKL5):c.2464del (p.Arg822fs) | VCV002947097 | 2947097 | 3105570 | | Pathogenic |
| NM_001323289.2(CDKL5):c.2469del (p.Glu824fs) | VCV000156652 | 156652 | 166506 | rs587783124 | Pathogenic |
| NM_001323289.2(CDKL5):c.2504del (p.Pro835fs) | VCV000143805 | 143805 | 153537 | rs267608660 | Pathogenic |
| NM_001323289.2(CDKL5):c.2522dup (p.Leu842fs) | VCV000280890 | 280890 | 264824 | rs886042014 | Pathogenic |
| NM_001323289.2(CDKL5):c.2529dup (p.His844fs) | VCV002945099 | 2945099 | 3108055 | | Pathogenic |
| NM_001323289.2(CDKL5):c.2529del (p.Leu843fs) | VCV000143806 | 143806 | 153538 | rs267608661 | Pathogenic |
| NM_001323289.2(CDKL5):c.2531dup (p.His844fs) | VCV000189582 | 189582 | 187626 | rs786204980 | Pathogenic |
| NM_001323289.2(CDKL5):c.2549del (p.Asn850fs) | VCV000156653 | 156653 | 166507 | rs587783125 | Pathogenic |
| NM_001323289.2(CDKL5):c.2572del (p.Arg858fs) | VCV000143807 | 143807 | 153539 | rs267608662 | Pathogenic |
| NM_001323289.2(CDKL5):c.2579_2582dup (p.Leu862fs) | VCV001075877 | 1075877 | 1065259 | rs2147178955 | Pathogenic |
| NM_001323289.2(CDKL5):c.2593_2616delinsG (p.Gln865fs) | VCV000451340 | 451340 | 446601 | rs1555955237 | Pathogenic |
| NM_001323289.2(CDKL5):c.2606del (p.Asn869fs) | VCV001070205 | 1070205 | 1065260 | rs2147178971 | Pathogenic |
| NM_001323289.2(CDKL5):c.2608dup (p.Ser870fs) | VCV000156654 | 156654 | 166508 | rs587783126 | Pathogenic |
| NM_001323289.2(CDKL5):c.2635_2636del (p.Leu879fs) | VCV000143809 | 143809 | 153541 | rs61753251 | Pathogenic/Likely pathogenic |
| NM_001323289.2(CDKL5):c.2636del (p.Leu879fs) | VCV001322054 | 1322054 | 1312372 | rs2147179000 | Pathogenic |
| NM_001323289.2(CDKL5):c.2673_2682del (p.Gln891fs) | VCV000870168 | 870168 | 858377 | rs1927142990 | Pathogenic |
| NM_001323289.2(CDKL5):c.2682dup (p.Pro895fs) | VCV000947226 | 947226 | 929639 | rs1569225454 | Pathogenic |
| NM_001323289.2(CDKL5):c.2682_2683insGGAA (p.Pro895fs) | VCV000573189 | 573189 | 574440 | rs1569225454 | Pathogenic |
| NM_001323289.2(CDKL5):c.2693_2694del (p.Arg898fs) | VCV001453219 | 1453219 | 1428505 | rs2147179071 | Pathogenic |
| NM_001323289.2(CDKL5):c.2821del (p.Tyr941fs) | VCV000643438 | 643438 | 649911 | rs1602300837 | Pathogenic |
